# A Mathematical Exploration of SDH-b Loss in Chromaffin Cells

**DOI:** 10.1101/2024.07.15.603520

**Authors:** Elías Vera-Sigüenza, Himani Rana, Ramin Nashebi, Ielyaas Cloete, Katarína Kl’učková, Fabian Spill, Daniel A. Tennant

## Abstract

The succinate dehydrogenase (SDH) is a four-subunit enzyme complex (SDH-a, SDH-b, SDH-c, and SDH-d) central to cell carbon metabolism. The SDH bridges the tricarboxylic acid cycle to the electron transport chain. A pathological loss of the SDH-b subunit leads to a cell-wide signalling cascade that shifts the cell’s metabolism into a pseudo-hypoxic state akin to the so-called Warburg effect (or aerobic glycolysis). This trait is a hallmark of phaeochromocytomas, a rare tumour arising from chromaffin cells; a type of cell that lies in the medulla of the adrenal gland.

In this study, we leverage the insights from a mathematical model constructed to underpin the metabolic implications of SDH-b dysfunction in phaeochromocytomas. We specifically investigate why chromaffin cells seemingly have the ability to maintain electron transport chain’s (ETC) Complex I function when confronted with the loss of the SDH-b subunit while other cells do not. Our simulations indicate that retention of Complex I is associated with cofactor oxidation, which enables cells to manage mitochondrial swelling and limit the reversal of the adenosine triphosphate (ATP) synthase, supporting cell fitness, without undergoing lysis. These results support previous hypotheses that point at mitochondrial proton leaks as a critical factor of future research. Moreover, the model asserts that control of the proton gradient across the mitochondrial inner membrane is rate-limiting upon fitness management of SDH-b deficient cells.

## 1. Introduction

The succinate dehydrogenase (SDH) is a four subunit complex (SDHa, SDH-b, SDH-c, or SDH-d) and an essential component of cell central carbon metabolism. It has two main functions: first, it accepts an electron from succinate to produce fumarate within the tricarboxylic acid (or TCA), and second, it catalyses the reduction of ubiquinone to ubiquinol as the mitochondrial electron transport chain (ETC) complex II (CII) [1, 2]. As such, it functions as a critical link between the TCA cycle and ETC [3, 1].

Many diseases and age related cell pathologies present SDH subunit mutations that cause partial to total loss of its functionality [2]. Needless to say, these metabolic disruptions result in catastrofic metabolic consequences [4, 5, 6]. However, the loss of its b subunit (SDH-b), in particular, is of special interest as it sets off a unique cascade of metabolic and signalling activities not seen in other mutations of this enzyme [4, 7, 8]. Dysfunction of the SHD-b subunit begins with a significant increase in intracellular succinate concentration [7, 8, 9, 10, 11]. Relative high succinate levels lead to inhibition of the 2-oxoglutarate (2-OG)-dependent hypoxia-inducible factor (HIF) prolyl-hydroxylases, stabilising HIF-2*α* in normoxic (or normal oxygen tensions) conditions leading to a pseudo-hypoxic state with metabolic consequences resembling those seen in cancer described as the ‘Warburg effect’ (or the so-called aerobic glycolysis) a hallmark of phaeochromocytomas [7, 8, 9, 10, 11]. These are rare tumours of the peripheral nervous system and adrenal glands, respectively [8, 9, 10, 12, 13].

Previous research has shown that in phaeochromocytomas, chromaffin cells (a type of endocrine cell located in the adrenal medulla in charge of producing and secreting catecholamines) preserve ETC complex I (CI) function despite the loss of SDH-b, unlike other cell types [7, 11, 14, 15, 16]. SDH-b defficient chromaffin cells appear to have a unique capactiy to remodel essential metabolic pathways and retain respiratory fitness under severe oxidative stress [11]. This adaptation is accompanied by considerable mitochondrial swelling, which is thought to be caused by the cell’s efforts to maintain a high membrane potential and ionic balance [17]. Furthermore, in SDH-b deficient chromaffin cells, the mitochondrial adenosine triphosphate (ATP) synthase (also known as ETC’s complex V) has been shown to function in reverse, hydrolysing rather than synthesising mitochondrial ATP to preserve membrane potential at the expense of cellular ATP [11, 18]. This is consistent with other mitochondrial diseases, in which it has been observed that blocking the mitochondrial ATP synthase can reverse its activity and restore cellular energy equilibrium [18, 19]. Similar findings in phaeochromocytomas have shed light on the link between SDH-b loss, mitochondrial enlargement, and CI functionality in SDH-b defficient chromaffin cells. However, the exact mechanisms underlying these events remain unclear [20, 21, 22, 23, 24].

In this study, we employ an in-silico dynamical model based on an immortalised mouse chromaffin cell (imCC) line model to investigate the metabolic consequences of SDH-b dysregulation in phaeochromocytoma [10, 11]. Our research builds upon the findings of Kl’učková et al. [11], who conducted extensive metabolic screenings on both wild type (WT) and imCC SDHb knockout (K.O.) variants to underpin the biology behind loss of SDH-b. Here we demonstrate how we develop a robust computational framework able to predict metabolic shifts and their cell wide consequences. We then characterise the causal mechanisms underlying the conclusions obtained by Kl’učková et al. [11]. Our main goal is to provide a first step towards a complete computational model able to explain why the loss of SDH-b activity, despite the enzyme’s widespread role in metabolism, predominantly impacts specific cell types such as chromaffin cells [11, 13, 25, 26].

## 2. Materials and Methods

### 2.1. Cell Culture and Chemicals

Previously characterised immortalised mouse chromaffin cell lines deficient in SDH-b (*SDH-b^−/−^ CL6 and CL8*) as well as their *SDH-b*^+^*^/^*^+^ counterparts were maintained in Dulbecco’s Modified Eagle Medium (DMEM) supplemented with 10% fetal bovine serum (FBS) and 1 mM pyruvate. All chemicals, including DMEM and FBS, were obtained from Sigma-Aldrich unless stated otherwise.

### 2.2. Total Cell Protein and Cell Growth Evaluation

1 *×* 10^6^ trypsinised cells were washed with PBS and lysed in 60 *µ*l of RIPA buffer for 30 minutes. Protein concentration in the cleared supernatant was determined using the BCA protein method (Thermo Fisher Scientific).

For cell growth measurements, cells were suspended in 200 *µ*l of Trypsin, followed by the addition of 400 *µ*l of PBS to achieve a total volume of 600 *µ*l. A 10 *µ*l sample was then pipetted into a cell counting grid chamber (Fast Read 102, Kova International). After loading the chamber with the sample, cells distributed across the chamber squares were counted. The grid consists of 10 squares, each with dimensions of 1 *×* 1 mm and a depth of 0.1 mm, corresponding to a volume of 0.1 *µ*l per square. The formula for determining the cell concentration (cells/ml) is given by:

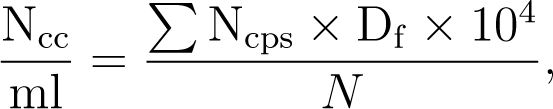

where, *N_cc_* is the number of cells counted, *N_cps_* is the number of cells per square, *D_f_* is the dilution factor, and N = 5 the number of squares in the grid. The dilution factor (*D_f_*) is 60 (Based on 600 *µ*l/10*µ*l, and the conversion factor = 10^4^ / 5.

### 2.3. Glucose, Lactate and Sodium measurements

Media was collected from each well, cells were spun down and supernatant was taken for analysis. Levels were measured using a Contour XT glucometer (Bayer).

### 2.4. Metabolic Tracing Experiments

For tracing experiments, cells were plated to be 70% confluent after 48 hours with 11 mM glucose and 2 mM glutamine, supplemented either in unlabelled form or as ^13^C_6_-glucose and ^13^C_5_-glutamine (CK Isotopes). After 48 hours, 100 *µ*l of media was removed for extraction and analysis. Cells were pelleted by centrifugation as described above. The remaining media was aspirated, and the empty wells were washed twice with ice-cold saline, after which 500 *µ*l of MeOH was added. Cells were scraped and transferred to a cold Eppendorf tube. Subsequently, 500 *µ*l of D_6_-glutaric acid in ice-cold water (1 *µ*g/mL) was added (CDN Isotopes, D-5227) followed by 500 *µ*l of chloroform (pre-chilled to -20*^◦^*C). After shaking on ice for 15 minutes and centrifugation, the polar phase was transferred to another tube for derivatisation, which was dried with centrifugation at 45*^◦^*C.

### 2.5. Derivatisation and Gas Chromatography – Mass Spectrometry

Dried down extracts were derivatised using a two-step protocol. Samples were first treated with 2% methoxamine in pyridine (40 *µ*l, or 20 *µ*l for primary samples, 1 hour at 60*^◦^*C), followed by the addition of N (tert butyldimethylsilyl)-N-methyl-trifluoroacetamide, with 1% tert - butyldimethyl - chlorosilane (60 *µ*l, or 30 *µ*l for primary samples, 1 hour at 60*^◦^*C). Samples were transferred to glass vials for GC-MS analysis using an Agilent 8890 GC and 5977B MSD system. One *µ*l of sample was injected in splitless mode with helium carrier gas at a flow rate of 1 mL per minute. The initial GC oven temperature was held at 100*^◦^*C for 1 minute before ramping to 160*^◦^*C at a rate of 10*^◦^*C per minute, followed by a ramp to 200*^◦^*C at a rate of 5*^◦^*C per minute, and a final ramp to 320*^◦^*C at a rate of 10*^◦^*C per minute with a 5-minute hold. Compound detection was carried out in scan mode. Total ion counts of each metabolite were normalised to the internal standard D_6_-glutaric acid.

### 2.6. Normalisation and Quantification

GC-MS data were analysed using Agilent Mass Hunter software for real-time analysis of data quality before conversion to .CDF format and analysis with in-house MATLAB scripts. Graphs and statistical analyses were performed using GraphPad Prism 9 and MATLAB.

### 2.7. Model Simulations and Code Availability

To partially quantify and parameterise the metabolic fluxes of our model, we employed ^13^C metabolic flux analysis (^13^C-MFA). Briefly, this method deduces intracellular flux patterns from mass isotopomer distributions measured via mass spectrometry. The process and established protocols were followed as described in Vera-Siguenza et al. [27], Antoniewicz [28], Young [29], Rahim et al. [30]. We conducted this analysis using the Isotopomer Network Compartmental Analysis (INCA) MATLAB routine, suitable for both steady-state and isotopically non-stationary metabolic flux analysis [29, 30].

All simulations and model code were executed using MATLAB and the ODE15s routine from MATLAB’s ODE suite. Source code for the models and figure generation can be freely accessed and obtained from our GitHub repository under an MIT open-source license [31, 32].

## 3. Model Construction

### 3.1. Assumptions

Our model consists of a system of ordinary differential equations. These monitor the rate of change in the concentrations of metabolites, critical ions, membrane potentials, and cellular and mitochondrial volumes with respect to time. Their rate of change is proportional to fluxes across four separate compartments: the extracellular space, cytoplasm, inner mitochondrial membrane space, and mitochondrial matrix, denoted by the subscripts *e*, *c*, *i*, and *m*, respectively (Fig. 1). Each flux in this model, *j_F lux_*, is described by a mathematical sub-model grounded in experimental observations or established mathematical concepts [33, 34, 35, 36, 37, 38].

**Figure 1:**
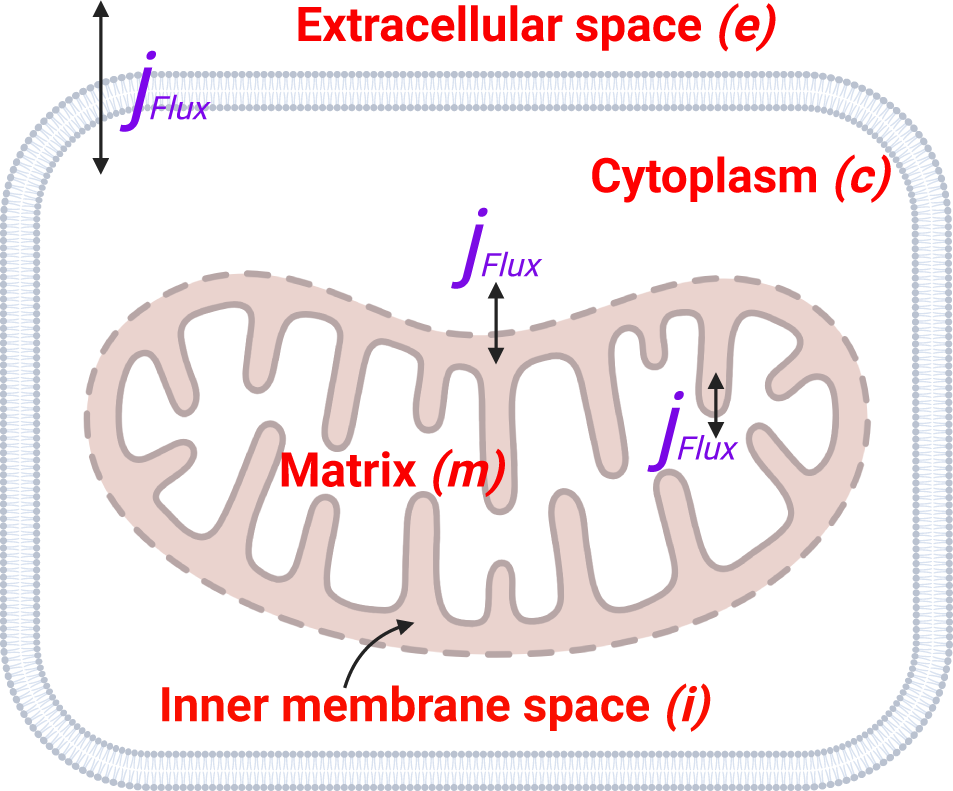
Schematic representation of the four compartments in the model: the extracellular space (*e*), cytoplasm (*c*), inner mitochondrial membrane space (*i*), and mitochondrial matrix (*m*). Each compartment is connected through various fluxes (*j_F lux_*) that describe the transport and transformation of metabolites and ions. The extracellular environment (*e*) is assumed to have constant metabolite and ion concentrations. (Figure created with BioRender.com [39])

Our model is based on mass conservation, with each metabolite concentration or ion species considered to be spatially uniform [36]. As a result, a concentration change in any of the chemical species occurs simultaneously throughout the compartment. Additionally, all extracellular metabolite and ion concentrations are kept constant this is evocative of a cell immersed in an infinite ion/metabolite solution. Although we recognise that this is a major simplification; for example, medium glucose and glutamine levels would decline with time, while lactate and other metabolites would increase during culture in-vivo, our modelling assumption is motivated by our need for computational efficiency and may more accurately represent the physiological situation with continuous flow of the interstitial fluid. Previous models, albeit making similar assumptions, have showed satisfactory predictive capacity [33, 34, 35]. Regardless of our assumptions, the model’s modularity enables for easy adaption to alternative configurations and modes in future studies, accounting for these dynamic changes as needed.

### 3.2. Ion channels and non-metabolism related fluxes

We adopted a model developed by Vera-Sigüenza et al. [33, 34, 35], which is based on the so-called Pump-Leak model [36, 40]. The ATP sodium/potassium (Na^+^/K^+^) pump (NaK-ATPase), is crucial to this paradigm, as it maintains cellular volume against osmotic pressures at the price of cellular ATP levels [36, 40, 41]. This process also requires the transmembrane movement of ions, specifically chloride Cl*^−^*, Na^+^, and K^+^, which is assisted by separate membrane ion channels and co-transporters or secondary active transport (see Supplementary Material S.1) [36].

Cl*^−^* regulation, is essential to maintain cellular osmotic balance. We equipped our model with a Cl*^−^*-Na^+^-K^+^ co-transporter (Nkcc) encoded by the Slc12a1/2 gene. In chromaffin cells, this co-transporter is crucial for maintaining elevated intracellular Cl*^−^* levels to activate Cl*^−^*-permeable GABA receptors [42]. This action leads to a depolarised chloride equilibrium potential. The mathematical model we used was adapted from the original works of Vera-Sigüenza et al. [33], Benjamin and Johnson [43], and later corrected by Palk et al. [44], and Gin et al. [45]. Furthermore, to adhere to the principle of mass balance, we introduced two generic efflux channels, one for Cl*^−^* and one for K^+^ (see Supplementary Material). These are modelled as simplified versions from those in Vera-Sigüenza et al. [33], resembling those in Keener and Sneyd [36] and Mori [40] (Fig. 2 see Supplementary Material S.2).

**Figure 2:**
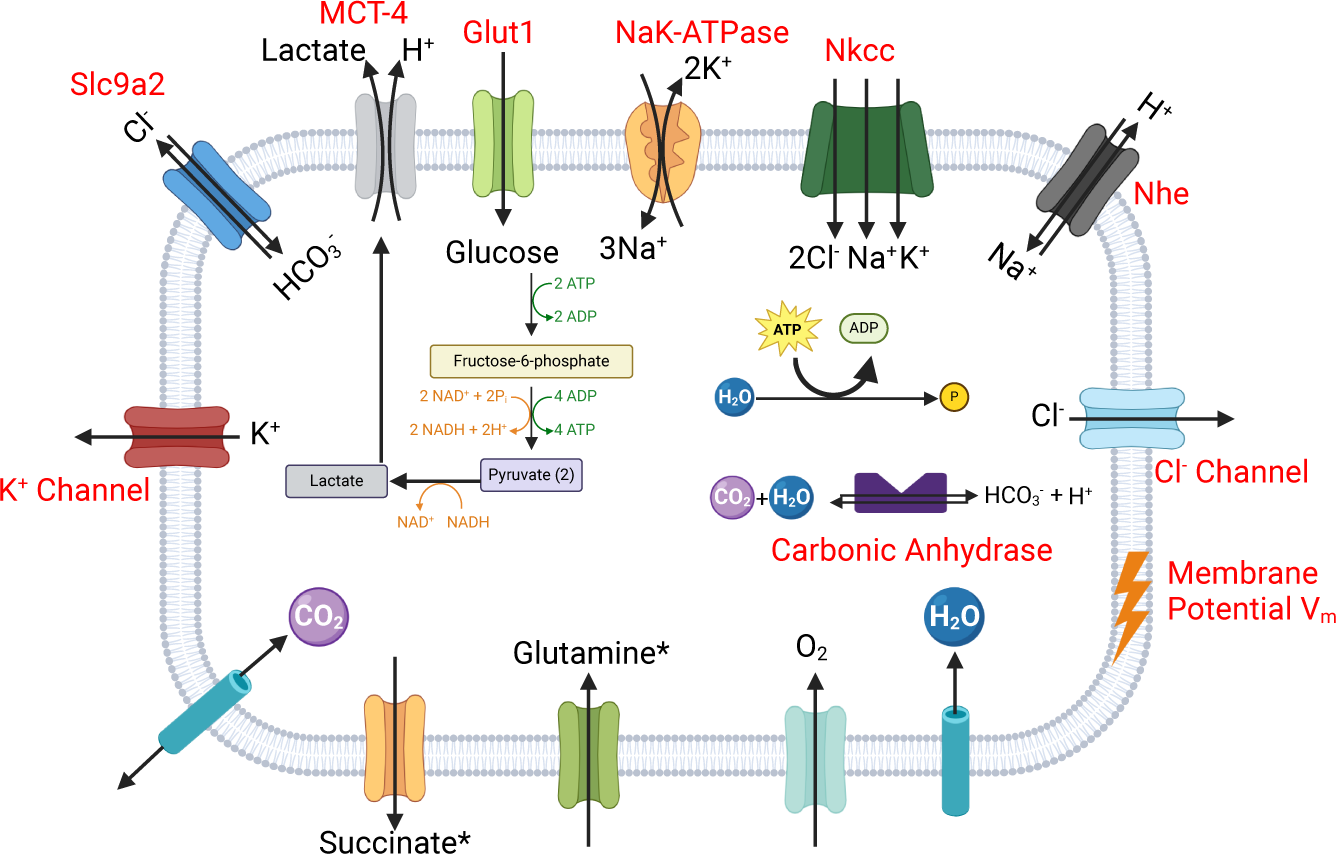
Schematic diagram of the cytoplasmic fluxes of the model. (Figure created with BioRender.com [39])

As stated above, the primary mechanism responsible for osmotic balance, the NaK-ATPase, maintains the necessary Na^+^ electrochemical gradient and energises all secondary active transports (i.e., ion transporters and co-transporters that rely on concentration gradients) in the model (Fig. 2) [36]. The submodel we employed (see Supplementary Material) consists of a simplified version of the mathematical construct developed by Crampin et al. [46] by Gin et al. [45] and later by Maclaren et al. [47]. Our approach accounts for the ATP dependency of the pump, as detailed in studies by Vera-Sigüenza et al. [33, 35] and Palk et al. [44] (see Supplementary Material S.3).

Our model also incorporates mechanisms for cellular pH regulation [48]. It includes a model for carbonic anhydrases essential for catalysing the interconversion between CO_2_ and water and the dissociated ions of carbonic acid HCO^+^ and H^+^ (see Supplementary Material S.4). The pH regulation system also includes the Na^+^-proton (H^+^) antiporter (see Supplementary Material S.5), and a lactate-H^+^ symporter (described as part of the glycolytic sub-model of our model as well as a succinate channel). As part of this pH regulatory system, we included a chloride-bicarbonate (HCO^+^) antiporter encoded by the Slc4a2 gene (see Supplementary Material S.6). This ubiquitous molecular machine enhances Cl*^−^* influx while expelling HCO^+^, following the principles outlined by Vera-Sigüenza et al. [35, 33] based on Falkenberg and Jakobsson [48] (Fig. 2).

The cytoplasmic volume in our model changes as a direct consequence of the osmotic gradient between its neighbouring compartments: extracellular space and mitochondrial matrix (see Fig. 1). In this model (see Supplementary Material S.7), the osmotic gradient between the cytoplasm and the extracellular space influences the cytoplasmic volume, while the gradient between the cytoplasm and the mitochondrial matrix regulates mitochondrial volume [27, 33, 34, 35, 36, 40, 49, 50]. Note, however, that this assumes that the plasma membrane cannot withstand hydrostatic pressure gradients. The mathematical submodel we employed is based on and adapted from Kedem and Katchalsky [35, 36, 44, 51].

Finally, to quantify the membrane potential (*V_m_*), established by the movements of charged ions across the cellular membrane, we employed Kirchhoff’s law via the so-called electric circuit model of the cell for a simple resistor-capacitor circuit (see Supplementary Material S.8) [33, 34, 35, 36, 40, 49]. This last addition assumes that the net sum of all currents in the circuit is zero. This has profound implications in our model as it essentially transforms the model to a system of differential-algebraic equations (i.e., a system of equations that contains both differential and algebraic equations) [40]. The full mathematical descriptions and equations for each of the mechanisms depicted in Figure 1 can be found in the Supplementary Material accompanying this article.

### 3.3. Glycolytic and TCA cycle model

Our glycolytic and tricarboxylic acid cycle (TCA) models are primarily based on the studies by Zhou et al. [37] and Salem [38] [52]. Briefly, these pathways facilitate the delivery of carbon, derived from glucose, directly into the mitochondrial compartment via the glycolytic pathway (see Supplementary Material S.9 S.11). This pathway encompasses ten enzymatic reactions and is localised in the cytosolic compartment of the cell (Figs.1 and 2). We included a sub-model that quantifies the reactions modulating the anabolism of pyruvate and lactate, and their subsequent incorporation into the cellular respiratory complex (Fig.3).

**Figure 3:**
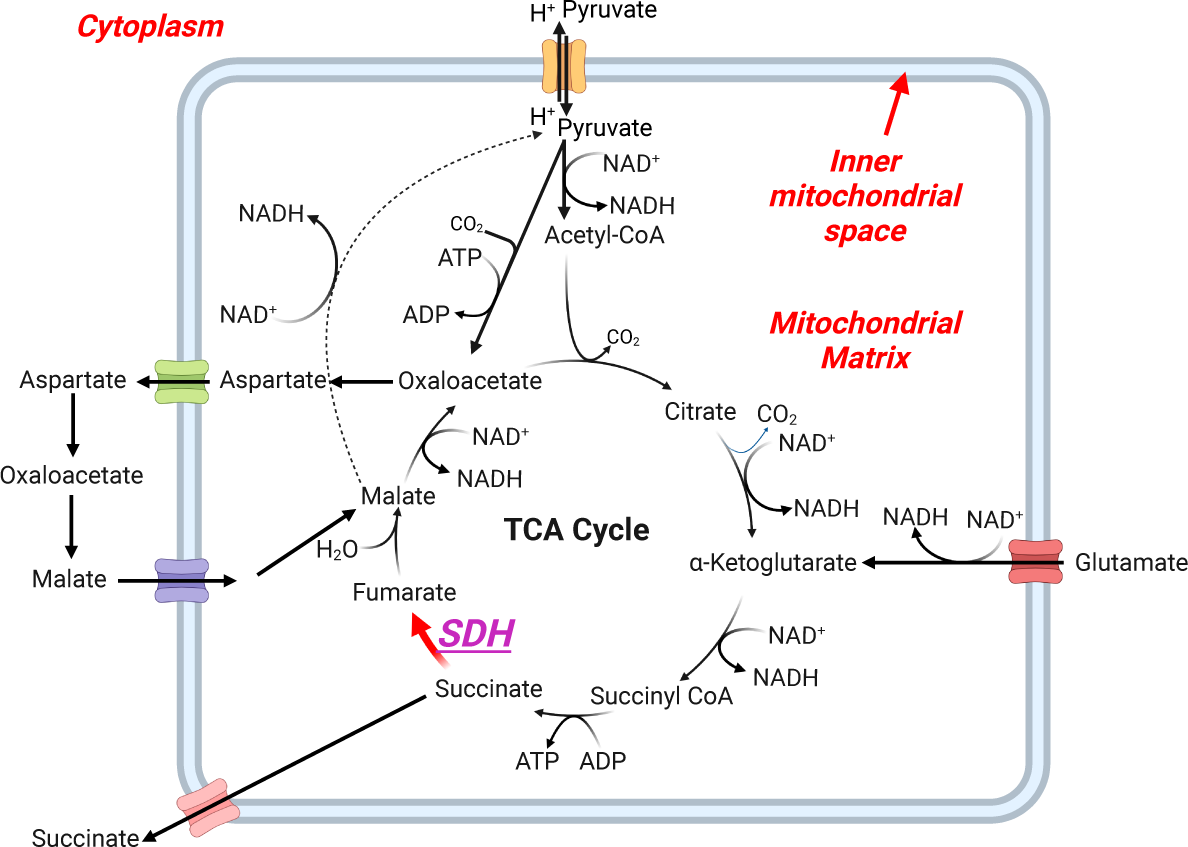
Schematic diagram of the mitochondrial matrix reaction fluxes of the model associated with the glycolytic and TCA cycle model. (Figure created with BioRender.com [39])

Unlike the models proposed by Zhou et al. [37] and Salem et al. [38, 52], we devised a net glycolysis reaction. Our simplification is driven primarily by the glucose influx facilitated by Glut1, a membrane glucose transporter encoded by the Slc2a1 gene and widely expressed in chromaffin cells, as well as by the production rate of pyruvate [53, 54]. Besides computational economy, this decision was motivated by our ability to obtain experimental data to accurately parameterise the entire pathway. This assumption allows us to focus on measurable values that can be directly obtained through experimental procedures, thereby allocating most of our computational resources to the electron transport chain. In other words, by aggregating the glycolytic reactions into a single net reaction, we streamline the model without compromising accuracy, focusing computational resources on solving the equations of the electron transport chain.

We acknowledge that this approach introduces potential model limitations, which we will address in the discussion section. However, our glycolytic model is able to calculate the rate of catalysis of carbon-source species as the difference between the rates of substrate production and utilisation, based on the availability of co-factors in the compartment. For our study purposes, this is sufficient [55]. The reaction flux sub-model is based on Michaelis-Menten kinetics and it incorporates the reaction flux between metabolic species and their corresponding reaction stoichiometric coefficients [11, 36, 56, 57]. Despite this, the model aligns well with experimental observations [37, 11].

The TCA equations take on the following form:

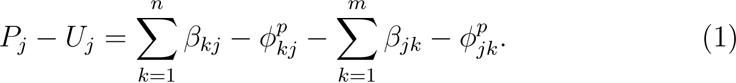

Here,

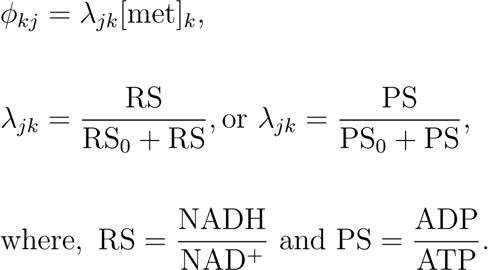

Eq. 1 details the production and utilisation of the *J^th^* metabolic species, which include all reactions resulting in anabolism or catabolism, are determined by the *n* reaction fluxes forming species *j* from species *k φ_kj_* and the *m* reaction fluxes forming species *k* from species *k φ_jk_*. The superscript *p* referes to the reaction processes in the TCA and *β_kj_* are the reaction stoichometric coefficents. The chemical reactions and their corresponding stoichiometries can be found in the supplementary data, a visualisation is provided in Fig. 3.

The rate coefficients *λ_jk_* are nonlinear functions of metabolite concentration ratios. These model the energetic state measured by ADP/ATP ratios, and the redox state by NADH/NAD^+^ ratios cytosolic and mitochondrial alike [38]. In this light, a particular ratio is only included in the rate coefficients of reactions where they participate as co-substrate or co-product. While these are Michaelis-Menten models [36], the concept of including these ratios has been successfully implemented and validated by Zhou et al. [37] and Salem et al. [52]. It is important to note that *λ_jk_* and*λ_kj_* denote the forward and reverse rate coefficients of a reversible reaction, respectively, and should not be defined separately in the context of reversible reactions.

Finally, we equipped the cytoplasm with the malate-aspartate shuttle reactions, which, together with lactate dehydrogenase, facilitate the redox conversion of NADH *↔* NAD^+^. This sub-model depends on two generic mitochondrial transporters: one for aspartate and one for malate (see Supplementary Material S.12)[36]. We term them ‘generic’ for two reasons: firstly, they are not the primary focus of detailed study within our model; secondly, we lack detailed knowledge of the specific transporters responsible for these metabolite fluxes, such as their dependence on calcium ions (Ca^2+^) [33, 35, 44, 49, 58]. Nonetheless, these transporters are essential to this study and to understanding the effects on central carbon metabolism in SDH-b deficient chromaffin cells [10]. Full mathematical descriptions and equations for each of the metabolic submodel mechanisms depicted in Figure 2 can be found in the Supplementary Material accompanying this article.

### 3.4. Equations of the electron transport chain model

The electron transport chain (ETC) comprises four enzymatic complexes. Complexes I, III, and IV function as proton (H^+^) pumps, moving H^+^s from the mitochondrial matrix to the inner membrane space (Fig. 4). These utilise electrons from NADH and ubiquinone reduction, as well as oxygen, to pump H^+^s against the established gradient. In contrast, complex II (or SDH) does not pump H^+^s but instead reduces ubiquinone for complexes III and IV. These molecular machines are coupled to ATP synthase, which uses the established electrochemical gradient (by the afford mentioned complexes) to pump H^+^s back into the mitochondrial matrix, driving ATP synthesis (Fig. 4).

**Figure 4:**
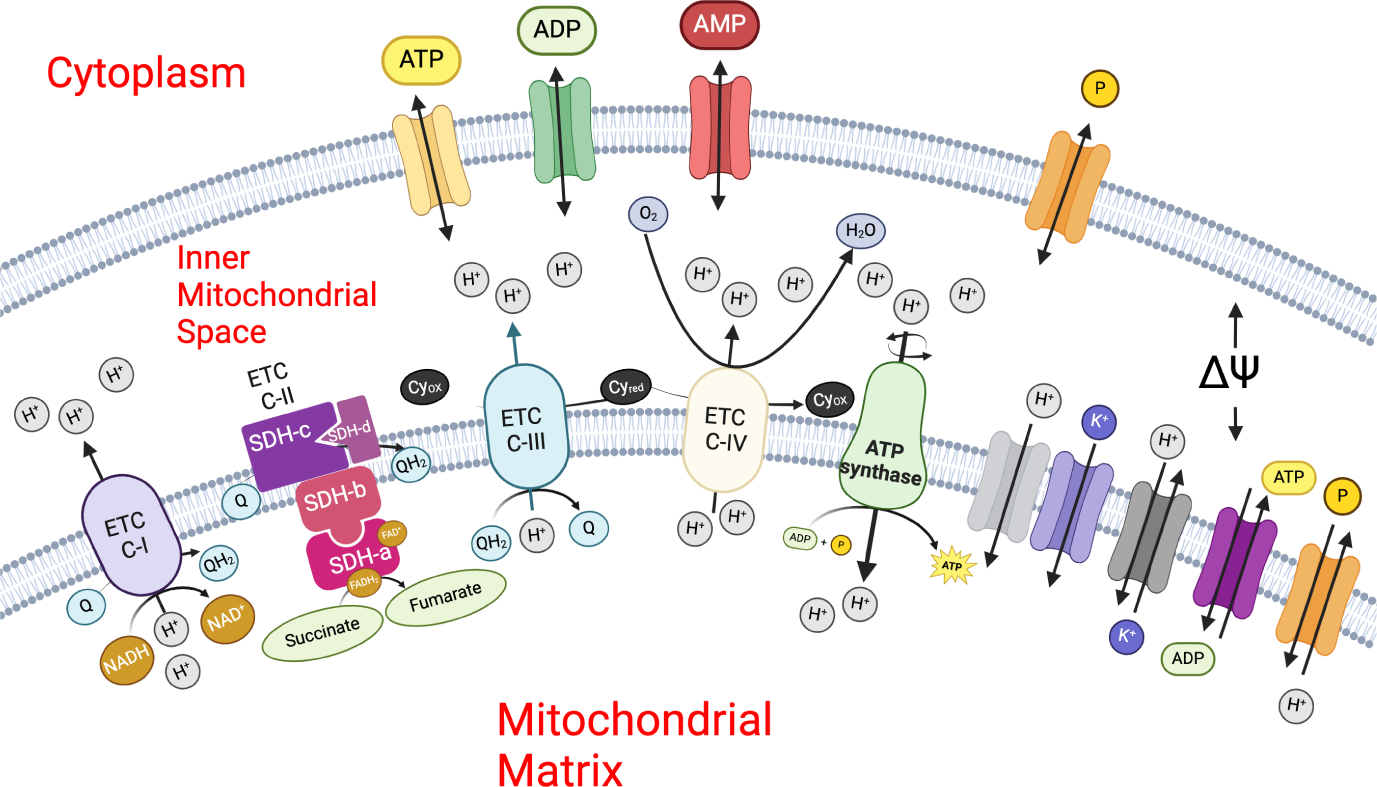
Schematic diagram of the inner mitochondrial membrane space reaction fluxes of the model associated with the electron transport chain model.

The mathematical construct included in our model is largely based on the models by Beard [59] and Manhas et al. [60]. However, we have modified some equations and parameters to ensure dimensional coherence with our data and the rest of the model. The conceptual model is briefly depicted in Fig. 4, with full details and equations provided in the Supplementary Material accompanying this article and the original works of Vera-Sigüenza et al. [34], Beard [59], and Manhas et al. [60].

#### 3.4.1. Proton motive force

The Proton Motive Force (PMF) represents the potential energy stored across a membrane. This force arises from the differential concentration of H^+^s on either side of the inner mitochondrial membrane (Fig. 4) [61]. The PMF is essential for cellular energy production, as it is utilised by the electron transport chain to actively transport protons from the matrix to the intermembrane space, thereby establishing both a concentration gradient and an electrochemical gradient, also known as the membrane potential difference.

We adapted a model of PMF by Beard [59] and modified it to include information from Johnson and Scarpa [61]. Briefly, the model relates the electrochemical gradient and the [H^+^] gradient across the inner mitochondrial membrane proportional to the difference in mitochondrial membrane potentials ΔΨ*_m_* (see Supplementary Material S.13). While the concentration gradient component reflects the variation in H^+^ concentrations between the intermembrane space and the mitochondrial matrix, in our model, the combined effect of these gradients creates the energy reservoir used by ATP synthase (Fig. 4).

#### 3.4.2. Electron transport chain complex I

Complex I (C1) is a H^+^ pump situated across the inner mitochondrial membrane (Fig. 4), initiates the electron transport chain by facilitating the transfer of electrons from NADH to ubiquinone (*Q*), reducing it to ubiquinol (*QH*_2_). This action energises the transport of H^+^ against the established chemical gradient from the mitochondrial matrix to the inner membrane space, hence contributing significantly to the mitochondrial PMF (see Supplementary Material S.14).

The enzymatic flux of Complex I is modelled as being proportional to the difference in concentrations of NADH and NAD^+^, as H^+^ fluxes drive the reaction. The flux is influenced by the free energy change (Gibbs’ free energy Δ*G* associated with H^+^ movement from one compartment to another.

#### 3.4.3. Electron transport chain complex III

Complex III mediates electron transfer from ubiquinol in the mitochondrial matrix to cytochrome c in the inner mitochondrial space (Fig. 4). Similar to Complex I, Complex III is a H^+^ pump and contributes to the mitochondrial PMF (see Supplementary Material S.15). The framework for this model is based on the design by Korzeniewski and Zoladz [62].

In this model, the flux of Complex III is defined by the influence of phosphate (*P_i_*) serves as a modulatory factor that drives respiratory activities to meet energy demands, a relationship first explored by Katz et al. [63]. Phosphate’s involvement underscores its significance in respiration or ATP synthesis reactions and highlights its potential impact on mitochondrial regulatory dynamics, including changes in volume.

#### 3.4.4. Electron transport chain complex IV

Similar to complexes I and III, Complex IV contributes to proton pumping (from mitochondrial matrix to inner mitochondrial space), which is instrumental in energising the ATP synthase. The model we use was first constructed by Korzeniewski and Zoladz [62] (see Supplementary Material S.16).

The flux through Complex IV depends on the oxygen concentration at any given time. This has profound consequences for the model kinetics, which vary non-linearly. Complex IV is also influenced by the proportion of reduced cytochrome c in the matrix relative to its total amount. As the concentration of reduced cytochrome c increases, so does the flux through Complex IV, given that reduced cytochrome c serves as a substrate for the reaction.

#### 3.4.5. Adenosine triphosphate synthase

The adenosine triphosphate (ATP) synthase plays a critical role in converting adenosine diphosphate (ADP) into ATP within the mitochondrial matrix. This model, constructed by Korzeniewski and Zoladz [62], centers around H^+^ movement across the mitochondrial membrane and how it energetically drives the synthesis of ATP (see Supplementary Material S.17). The process depends on the concentration gradients between ADP and ATP, along with phosphate (*P_i_*) in the mitochondrial matrix, ensuring the reaction is energetically favourable.

#### 3.4.6. Adenine nucleotide translocator (ANT)

The adenine nucleotide translocator (ANT) flux plays a critical role in cellular energy management by facilitating the displacement of one negative charge from the mitochondrial matrix to the mitochondrial inner membrane space. In our model (see Supplementary Material S.18), this process is coupled to the electrostatic membrane potential and modelled as a membrane transporter according to Halestrap and Brenner [64], Korzeniewski and Zoladz [62], Korzeniewski [65] and Keener and Sneyd [36].

#### 3.4.7. Magnesium-ATP binding

Magnesium (Mg^2+^) plays a crucial role in the stability and function of ATP, which comprises a ribose sugar, adenine, and three negatively charged phosphate groups. By forming bonds with these phosphate groups, magnesium effectively reduces their inherent repulsion [66, 67]. This interaction (see Supplementary Material S.19) is essential for enzymes such as kinases that rely on ATP-Mg^2+^ complexes to enhance catalytic efficiency. The inclusion of Mg^2+^ in our model coordinates the phosphates during ATP hydrolysis. It also serves as a preamble for a future exploration of the metabolic/signalling intersectionality.

#### 3.4.8. The mitochondrial phosphate carrier (PiC Slc25a3)

The PiC, encoded by the solute carrier family 25a3 (Slc25a3), is a crucial protein within the mitochondria responsible for transporting phosphate across the inner mitochondrial membrane [58, 68]. In our model, it facilitates the movement of inorganic phosphate between the matrix and the mitochondrial intermembrane space, coupled to the H^+^ gradient (see Supplementary Material S.20).

The transport mechanism model, based on that by Korzeniewski and Zoladz [62], involves a cotransport process where H^+^ and dihydrogen phosphate (H_2_PO_4_) are moved together in a 1:1 ratio, allowing for an electroneutral exchange across the membrane. The association of H^+^ with *P_i_* is in equilibrium, effectively balancing the phosphate species on both sides of the membrane.

#### 3.4.9. Adenyl kinase

Adenylate kinase (AK) is an essential enzyme for maintaining cellular adenine nucleotide balance [69, 70]. In the mitochondrial inner membrane space, AK catalyses the transfer of high-energy phosphates among ATP, ADP, and AMP, a process vital for cellular energy management. This reaction is modelled to proceed via a general linear equation, applicable across different isozymes (see Supplementary Material S.21). The model incorporates an equilibrium constant, XAK, and an enzyme activity parameter, XAK, to quantify the flux of nucleotides mediated by AK, effectively describing its role in the energetic equilibrium within cells.

#### 3.4.10. Proton leak and potassium fluxes

In this study, we assume that the effects of Ca^2+^ concentrations and fluxes on membrane potential are secondary compared to the respiratory chain, adenine nucleotide translocator (ANT) current, and proton leaks. Consequently, Ca^2+^ fluxes are not incorporated at this stage but are planned for inclusion in future iterations of the model. However, K^+^ and magnesium (Mg^2+^) ions are integral to the model due to their roles in buffering matrix pH and facilitating ATP synthesis and ANT flux, respectively. The movements of K^+^ and H^+^ across the mitochondrial membrane are modelled using the Goldman-Katz-Hutchkin equation, a solution derived from the onedimensional Nernst–Planck equation [36] (see Supplementary Material S.22).

#### 3.4.11. Potassium/Proton exchanger

The mitochondrial potassium/proton exchanger is vital for the transport of K^+^ into the mitochondrial matrix in exchange for H^+^. This exchanger is critical in maintaining the mitochondrial membrane potential and the pH gradient, both essential for ATP production via oxidative phosphorylation [71]. In our model, the dynamics of this exchanger are captured using a linear exchange model as outlined by Keener and Sneyd [36] (see Supplementary Material S.23).

#### 3.4.12. AMP, ADP, and ATP

Transport of ATP, ADP, AMP, and Pi between the cytosol and the mitochondrial inter-membrane space is modelled using linear transfer between compartments (see Supplementary Material S.24).

#### 3.4.13. Succinate dehydrogenase/electron transport chain complex II

Our model is an adaptation from a model first constructed by Manhas et al. [60]. At the core of this model are the binding polynomials that describe the likelihood of various molecules binding to specific sites on the SDH. These polynomials arise as the denominator of a rational function that represents the average number of occupied binding sites as a function of ligand (substrate) activity [36]. This approach enables the capture of the mechanistic dynamic interaction of substrates and inhibitors with the enzyme, providing a quantitative measure of binding affinities and their impacts on its activity. The specific formulations of these binding polynomials highlight the interactions of ligands such as ubiquinone within the SDH (see Supplementary Material S.25).

Additionally, the model adjusts the redox potentials of various SDH-associated redox centres to account for pH variations, which can significantly impact the enzyme’s electron transfer capabilities (see Supplementary Material S.26). These corrections ensure that the model is able to attain physiological conditions more accurately, allowing for a better understanding of how pH shifts influence the redox state of the enzyme [60].

The fluxes associated with SDH, including the transfer of electrons from succinates through various bound states (or SDH-b subunit) (see Supplementary Material S.27) to the eventual reduction of ubiquinone (SDH-c and SDH-d), are based on the established binding and redox potential models. This comprehensive model ensures that all critical aspects of SDH functionality, from substrate binding to electron transfer, are captured with high fidelity.

### 3.5. Model Equations

The full model equations can be found in the Supplementary Material S.28. Moreover, given the extensive list of parameters involved in this model we have provided, as part of the supplementary data, two files that contain all the parameters and their values see Supplementary Material.

## 4. Experimental data integration and model parameter fitting

To elucidate the complex metabolic interplay within chromaffin cells, we conducted an isotopic ^13^C labelling experiment. This method traces carbon atoms through key metabolic pathways, providing insights into substrate consumption, intermediate metabolite dynamics, and end-product generation [28]. Our main purpose was to quantify fluxes, parameterise reactions, refine our model, and validate our findings against experimental work [11, 72, 8]. For this purpose, we utilised ^13^C_6_-glucose (uniformly labelled glucose) due to its proven ability to comprehensively reveal carbon utilisation in central carbon metabolic processes, as assessed by gas chromatography-mass spectrometry (GC-MS) [28, 29]. This technique, combined with ^13^C-metabolic flux analysis (^13^C-MFA), enabled us to dissect and compare the metabolic fluxes between wild-type (WT) and SDH-b knockout (K.O.) imCC cells [28].

### 4.1. Isotopic labelling

Following the protocols outlined by Antoniewicz [28] and Vera-Siguenza et al. [27], we began by monitoring the proliferation rates of WT and SDH-b K.O. cell lines over a 48-hour period (Fig. 5a). We observed that SDH-b K.O. cells had a significantly lower growth rate (0.021756 h*^−^*^1^) compared to WT cells (0.034426 h*^−^*^1^). This indicates a substantial reduction in growth for cells lacking SDH-b, aligning with findings from previous studies [11, 8]. The growth rates were calculated by assuming cells continuously divide [28]. Thus, we expect exponential growth according to:

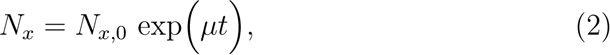

**Figure 5:**
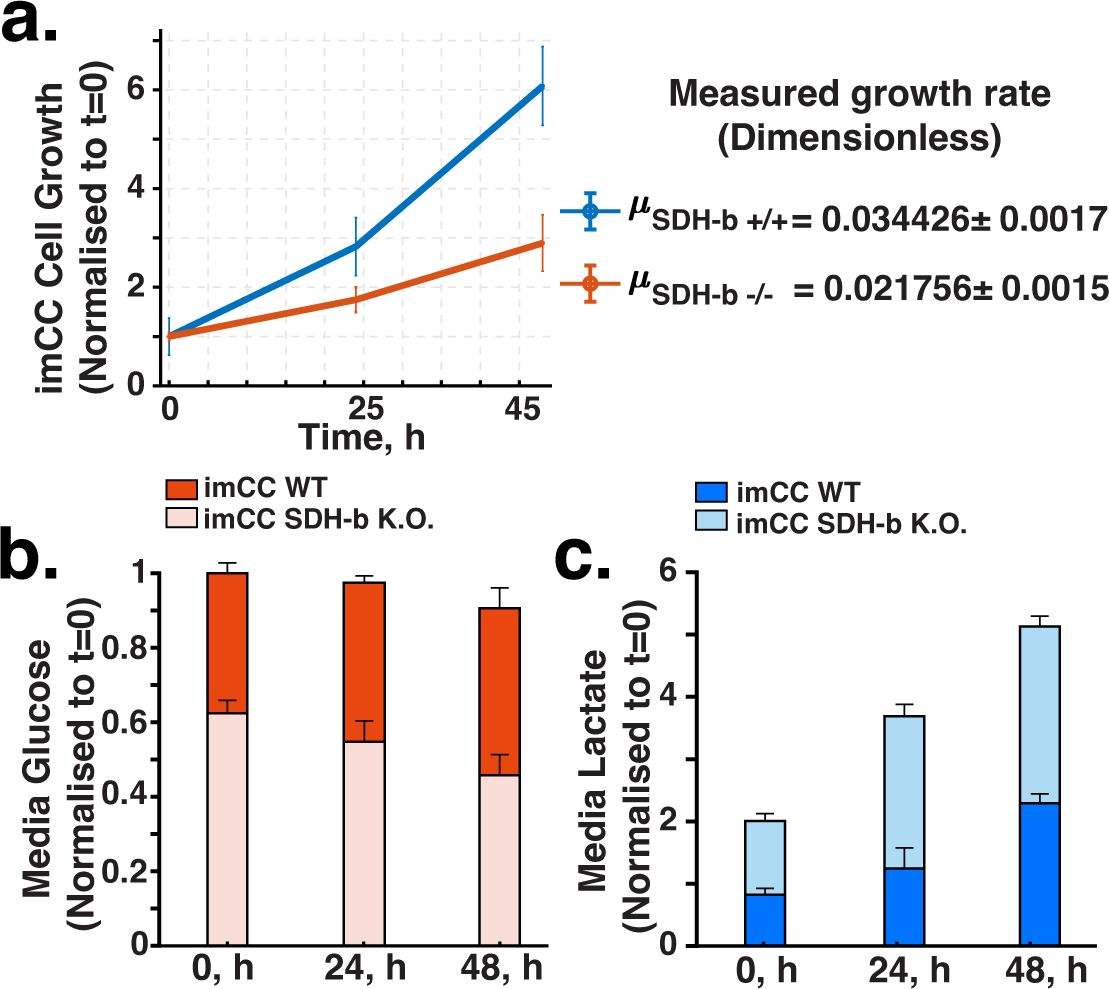
Extracellular and growth measurements. a. WT vs. SDH-b K.O. growth rate monitored over a 48 hour period. Rate parameters were obtained assuming exponential doubling time. b. WT vs. SDH-b K.O. glucose consumption comparison. SDH-b K.O. cells show a decrease in consumption due to impared TCA cycle. b. WT vs. SDH-b K.O. lactate efflux comparison. SDH-b K.O. cells show a marked increase in lactate efflux. This is indicative of a shift towards aerobic glycolysis. (For absolute concentration values in figures b. and c. refer to Supplementary material.)

where *N_x_*represents the number of cells and *µ* is the growth rate (1/hr).

Solving for *µ*, we obtain:

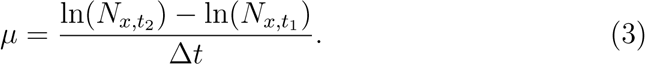

Subsequently, we assessed glucose and lactate levels in the media to gauge metabolic activity. Over the same period, both WT and SDH-b K.O. cells demonstrated distinct glucose consumption patterns. We noticed that SDHb K.O. exhibited reduced glucose intake compared to WT cells (Fig.5b).

However, SDH-b K.O. cells exhibited an increased lactate production rate, corroborating the expected shift towards aerobic glycolysis, a phenomenon typically observed in these cells (Fig.5c) [13, 23, 10, 7, 8]. Using these measurements, we were able to calculate the glucose and lactate external rates:

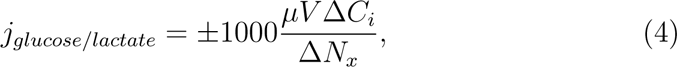

here Δ*C_i_* (mmol/L) is the change in concentration of the *i^th^* metabolite between two sampling time points, Δ*N_x_* is the change in cell number during the same time period, V (mL) is the culture volume (see Materials and Methods), and *µ* (1/h) is the growth rate [28]. Note that in eq. 4, consumption is defined negative and metabolite secretion is defined postive.

Following preliminary assessments, we undertook comprehensive isotopic labelling using ^13^C_6_-glucose (details in Materials and Methods). We then analysed the isotopic labelling patterns of key tricarboxylic acid (TCA) cycle metabolites, including lactate, citrate, succinate, malate, fumarate, and aspartate (Fig.6a). All simulations in our ^13^C-MFA underwent sum of square residuals statistical test (SSR aims at recovering a sparse weight vector in an underdetermined linear model with a known and fixed dictionary matrix) to assess the goodness-of-fit, parameter confidence intervals, and model identifiability evaluation (Fig.6b). This enhanced the reliability of our findings [29]. The mass isotopomer distributions (MIDs) for these metabolites revealed distinct differences between SDH-b K.O. and WT cells, suggesting a potential rewiring of the TCA cycle in SDH-b deficient cells (Fig. 6a and Supplementary Materials).

**Figure 6:**
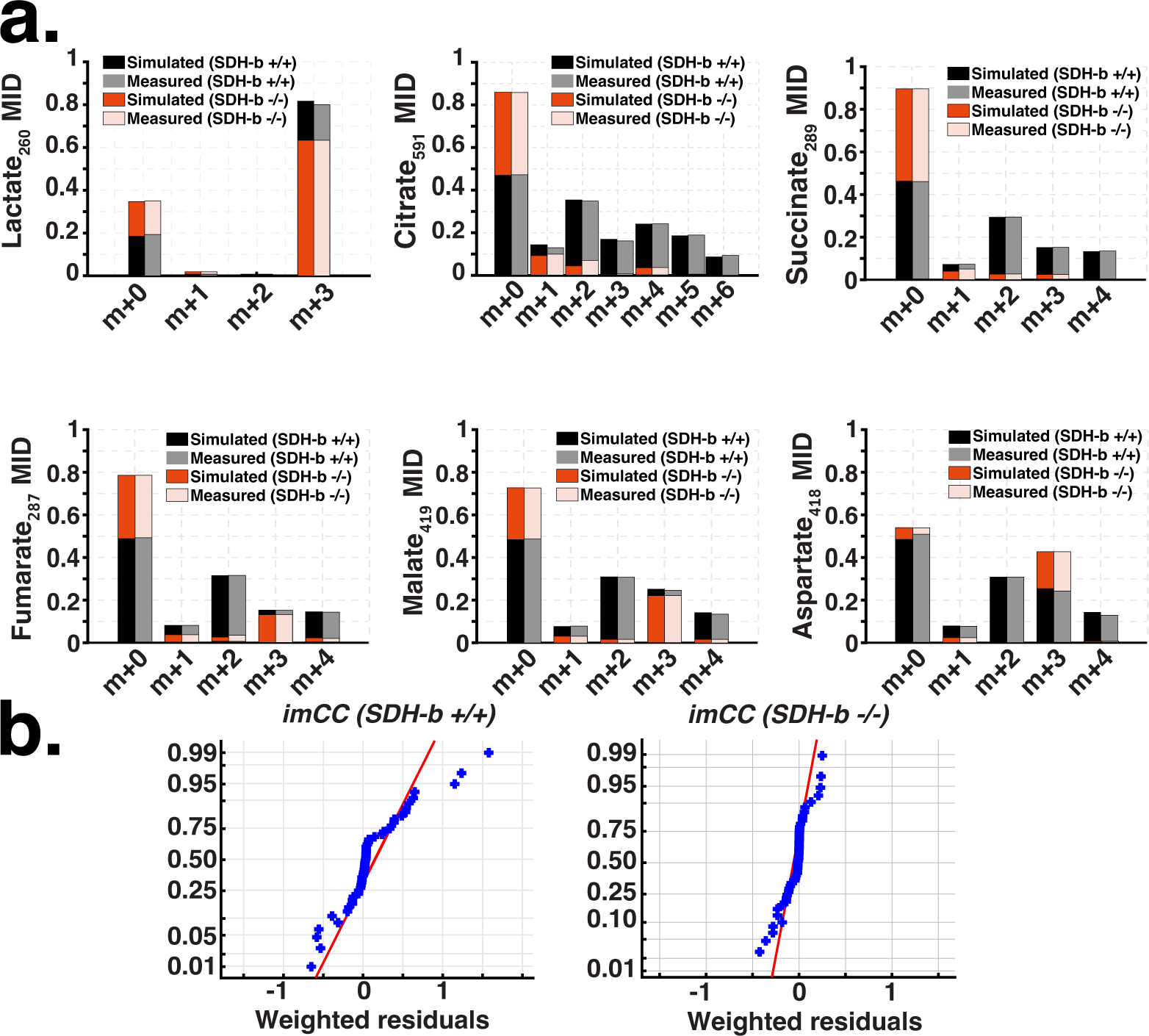
Experimental vs. simulation mass isotopomer distributions from ^13^C_6_glucose labelling experiment (WT vs. SDH-b K.O.). a. ^13^C-MFA predicted mass isotopomer distributions of several metabolites across the central carbon metabolism pathways. b. Assesment of good-fit using sum of squares due to error (SSR), or the quantity which the least squares procedure attempts to minimise to attain the observed mass isotopomer distributions. The results obtained provide a good fit to experimental data.(see Suplementary Material)

We then performed ^13^C-metabolic flux analysis (^13^C-MFA) to obtain the fluxes for the central carbon metabolism reactions (see Supplementary Material). Our analysis revealed that fluxes from glucose to pyruvate remained consistent across WT and SDH-b K.O. cells, indicating unaltered glycolytic rates regardless of SDH-b status (Fig. 7a & b). Nevertheless, an increase in lactate production in SDH-b K.O. cells signalled enhanced lactate dehydrogenase activity, which is likely to be compensation compensating for TCA cycle disturbances resultant from SDH-b loss [41, 7, 8].

**Figure 7:**
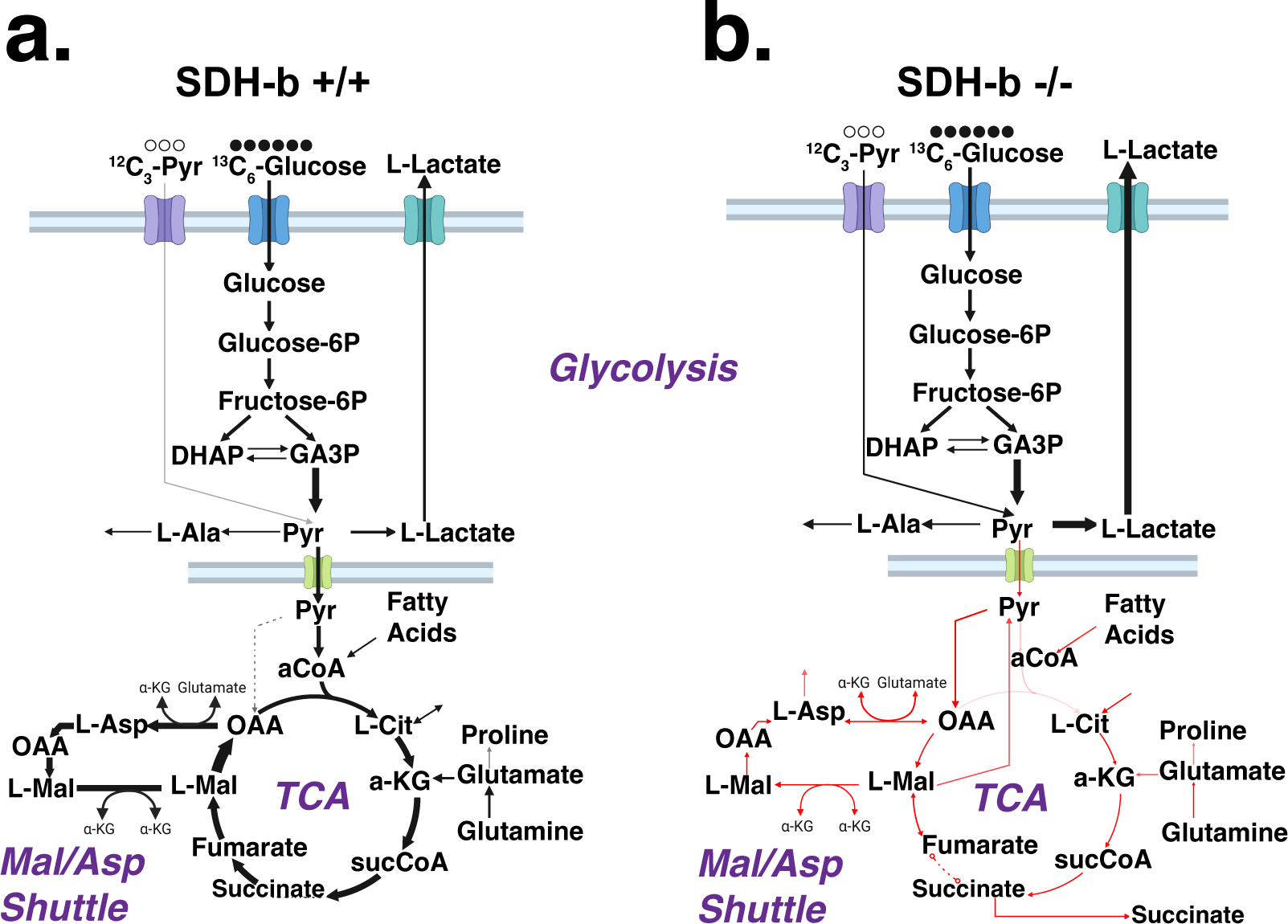
Comparison of ^13^C_6_-glucose labelling experiment-derived fluxes (metabolic flux analysis). a. imCC WT flux map. b. imCC SDH-b K.O. flux map. The flux alterations in succinate-related reactions suggest a bottleneck, prompting cells to employ compensatory mechanisms such as a reversal of the malate-aspartate shuttle and increased reliance on pyruvate carboxylase over pyruvate dehydrogenase.

Our simulations also indicate that SDH-b K.O. cells exhibit significantly altered fluxes in TCA intermediate steps, involving succinate, directly attributable to SDH-b loss (Fig. 7b). The flux alterations in succinate-related reactions indicate a bottleneck, prompting cells to employ compensatory mechanisms such as a decoupling of the malate-aspartate shuttle and increased reliance on pyruvate carboxylase over pyruvate dehydrogenase. A similar outcome was reported by [16]. As a consequence, our analysis suggests, a possible increased reliance on exogenous fatty acid pathways could be responsible for sustaining the loss of pyruvate dehydrogenase activity, which directly affects the citrate synthesis arm of the TCA cycle (e.g., lipid metabolism, fatty acid *β*-oxidation and oxidative phosphorylation) [7, 8, 10].

### 4.2. Enrichment and rate parameters

After obtaining the fluxes of the model (Fig. 7) through ^13^C metabolic flux analysis (MFA), we proceeded to carry out a simulation of isotopic labelling dynamics, or enrichment measurements [28]. Briefly, an isotopic enrichment measurement is a technique used to quantify the relative abundance of different isotopes of an element in a sample. For example, in metabolic studies, a sample might be enriched with ^13^C-glucose. By measuring the ^13^C/^12^C ratio in different metabolites, it is possible to track the incorporation and flux of carbon through metabolic pathways, as well as enzymatic reaction rates [73].

After obtaining the fluxes of the model (Fig. 7) through ^13^C -Metabolic Flux Analysis (MFA), we proceeded to carry out an enrichment simulation [28]. This approach stems from the realisation that the resulting fluxomic data implicitly provides the rates at which enzymes involved in central carbon metabolism catalyse their respective reactions. Carrying out an enrichment simulation, allowed us to determine key kinetic parameters from our own dataset. The result is that, effectively, the model recapitulates more accurately the phyisiological traits of the chromaffin cell model (imCC) [29].

To achieve this, and given that the majority of our model central carbon metabolism pathways primarily follow Michaelis-Menten kinetics, we sought the *V_max_* and respective *K_d_* parameters [36]. However, enrichment percentages derived from ^13^C-labelling experiments do not directly equate to concentration measurements; this is an issue as our model deals with changes in concentration over time.

To overcome this, we averaged the enrichment of the isotopomers and normalised it to *V_max_*= 1. Using this approach, we set *V_max_* = 1 for our model equations and then deduced the corresponding *K_d_* and Hill coefficients to align with the rates of enrichment. In this frame, we are able to deduce the rate of catalysis of the different enzymes in our model. Once the value was obtained, we then applied a non-linear optimisation MATLAB routine (optimisation toolbox’s Levenberg-Marquardt see Supplementary Material) to deduce the *V_max_*parameter given we know metabolite concentrations (from raw ion counts see Supplementary data), *K_d_*, and flux from our ^13^C-MFA simulation.

Thus, to re-iterate, the rationale for equating rate parameters with enrichment rates is as follows: enrichment rates, which indicate the proportion of molecules in a metabolite pool labelled with ^13^C, serve as an indirect measure of metabolic flux through a pathway. As flux increases, so does the proportion of labelled molecules, analogous to how *V_max_* reflects the maximal catalytic capacity of enzymes, thereby dictating the rate of metabolic reactions. By aligning the Hill function parameters with the enrichment rates, we capture the interplay between substrate availability and enzyme activity, essentially allowing the Hill function parameters particularly *V_max_* and *K_d_* to act as proxies for enrichment rates.

In Figures 8 and 9, it is possible to see the results of these simulations and parameter estimation. Here, the enrichment percentages reflect the proportion of metabolites containing the ^13^C label, indicating the flux through metabolic pathways that incorporate ^13^C from our tracer into the pathway of the metabolite of interest. Fig. 8a and b illustrate the enrichment curve for pyruvate over time, with a model fit based on a glycolysis model. The curve also depicts normalised enrichment over time, with the x-axis representing time in hours and the y-axis representing normalised enrichment. Similar to 8b, the model fit aligns closely with the measured data, demonstrating the effectiveness of the glycolysis model in capturing pyruvate enrichment.

**Figure 8:**
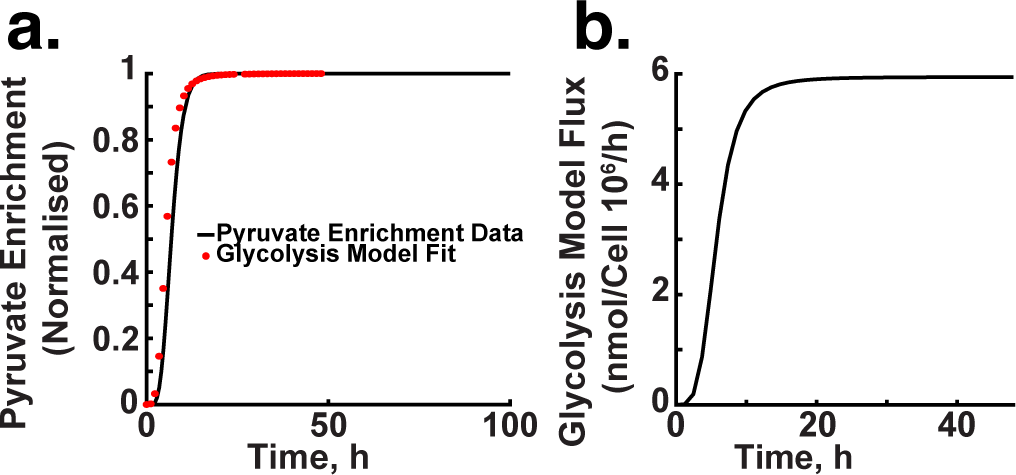
Enrichment simulation. a. and b. display the mean enrichment curve of pyruvate over time, aligned with a glycolysis model. The graphs show the model’s fit to the normalised enrichment data, highlighting the model’s accuracy in representing pyruvate dynamics in glycolysis.

**Figure 9:**
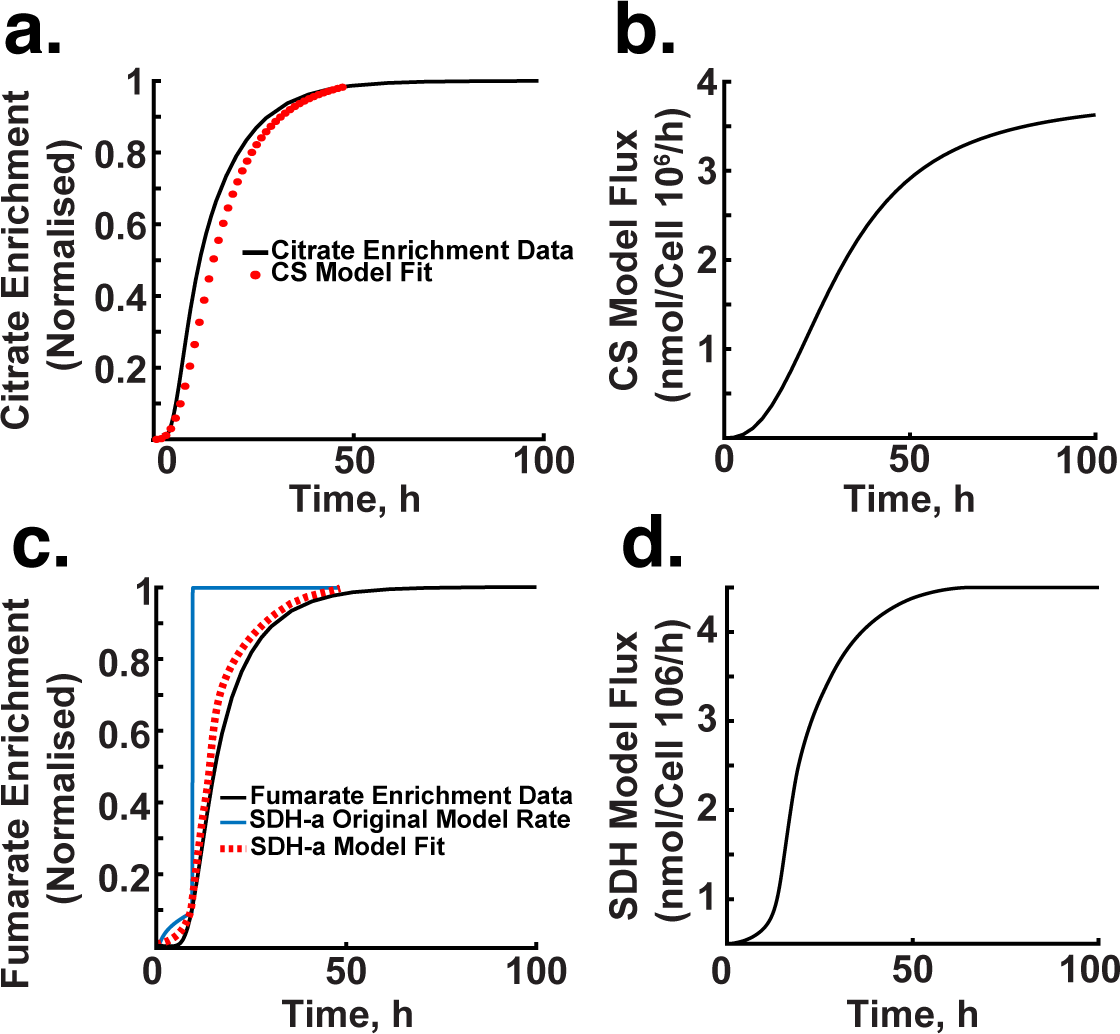
Enrichment simulation. a. and b. depict how enzymatic activity affects the production of citrate over time setting the incorporation of carbons into the TCA, with the curve indicating the peak rate of citrate synthesis by citrate synthase under optimal substrate availability. c. and d. Depict the enrichment curve for fumarate, including both the original and adjusted model fits, highlighting the adjustments made to the SDH model parameters for better alignment with experimental data. panel d. shows the estimated flux over time using the SDH-a model (imCC WT).

We also used the enrichment data to describe how the enzymatic activity leading to labelled citrate production responds over time (Fig 9a and b). In this context,*V_max_* represents the maximum proportion of ^13^C-labelled citrate production when substrate (acetyl-CoA and oxaloacetate) availability is at its highest and most effective for the enzyme citrate synthase. Notice how the incorporation of carbons into the TCA cycle takes a substantially longer amount of time compared to the rate of pyruvate anabolism by the glycolytic pathway.

Fig.9c shows the enrichment curve for fumarate over time. This panel also includes the original model rate and the fit obtained through parameter adjustments of the SDH model. The original model by Manhas et al. [60] rate deviates significantly from the measured enrichment data. We adjusted model parameters to fit and align closely with the data, demonstrating the effectiveness of parameter manipulation in achieving a more accurate representation of SDH activity. Fig.9d depicts our estimation of flux of the SDH-a model (imCC WT) over time.

We repeated the simulations (see Supplementary material) for all the measured metabolites in our labelling experiment. Thus, we were able to characterise the glycolytic and TCA cycle metabolic sub-models using this approach.

## 5. Model results

### 5.1. Model benchmarking

Having attained model parameterisation, we benchmarked the model to ensure its physiological relevance. This was carried out to ensure the model could reliably simulate known biological behaviours, thereby confirming its utility in predicting metabolic responses under well known conditions [74].

As such, we began by evaluating the model’s response to oxygen depravation (or hypoxia). Essentially, we simulated conditions where oxygen levels were decreased. The model, in turn, should display known significant metabolic shifts (e.g., changes in electron transport chain activity, alterations in ATP production, and variations in metabolite fluxes) [75, 76].

The simulation results depicted in Figure 10 align closely with established physiological responses to hypoxia [7, 77, 78]. In Fig. 10a it is possible to see the model’s response to a decrease in oxygen flux leads to a reduction in the flux through the mitochondrial pyruvate carrier. This is consistent with the known decrease in mitochondrial oxidative metabolism under low oxygen conditions [79, 80]. The model indicates that this adaptation helps conserve oxygen by reducing its utilisation in the mitochondria.

**Figure 10:**
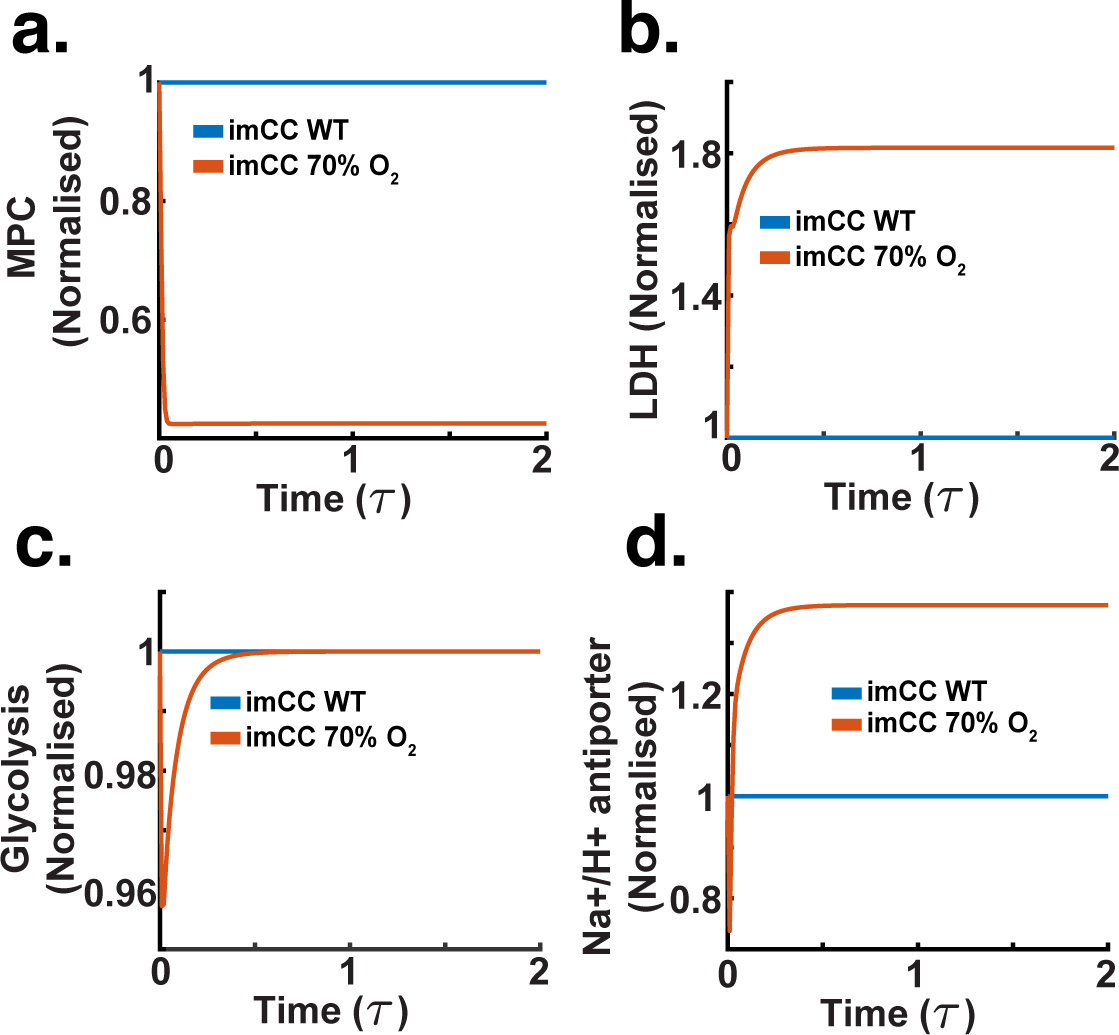
Benchmarking simulation (dimensionless time *τ*). We subjected the chromaffin cell model to hypoxia (a 30% reduction in oxygen flux). a. Relative activity of the mitochondrial pyruvate carrier, showing a decrease as hypoxia progresses over time. b. Relative activity of lactate dehydrogenase, illustrating an increase due to enhanced anaerobic glycolysis. c. Relative glycolysis rate, maintaining steady activity despite hypoxic conditions.d. Relative activity of the (Na^+^/H^+^) NHE antiporter. The antiporter increases its activity to remove H^+^s from the cytosol, protecting cells from acidification as a result of anaerobic metabolism.

Fig.10b depicts an increase in lactate dehydrogenase activity a welldocumented reaction to hypoxia, where cells increase lactate production to regenerate NAD^+^ for continued glycolytic ATP production [9]. In Fig.10c, a slight upregulation in glycolysis activity further supports this metabolic shift, indicating an increased reliance on glycolytic pathways to meet energy demands when oxidative phosphorylation is compromised. Additionally, Fig. 10d demonstrates an increase in the activity of the Na^+^/H^+^ antiporter, indicative of the cellular effort to regulate intracellular pH under acidic stress conditions commonly induced by increased lactate production. This response is critical, as the ability of the NHE to regulate acidity changes in the cytoplasm signifies the model’s correct response to hypoxia [81, 27].

The model also highlighted significant changes in intracellular metabolite concentrations and pH, reflecting typical cellular adaptations to hypoxia. Fig.11a illustrates a decrease in intracellular pH, which is a consequence of the accumulation of acidic metabolic by-products such as lactate. This acidification is a hallmark of the hypoxic response, often leading to the activation of pH-regulating mechanisms such as the Na^+^/H^+^ antiporter observed in Fig.10d. Fig.11b shows an increase in intracellular NADH concentration, which is expected due to the reduced activity of the electron transport chain under low oxygen conditions, causing an accumulation of reduced cofactors [78]. Fig.11c shows a rise in intracellular lactate concentration, further supporting the increased glycolytic activity and lactate production, while Fig. 11d indicates a reduction in intracellular pyruvate concentration, reflecting its increased conversion to lactate. These shifts in metabolite levels are characteristic of the metabolic shift that occurs during hypoxia, aiming to adapt to reduced oxygen availability while maintaining cellular energy balance.

**Figure 11:**
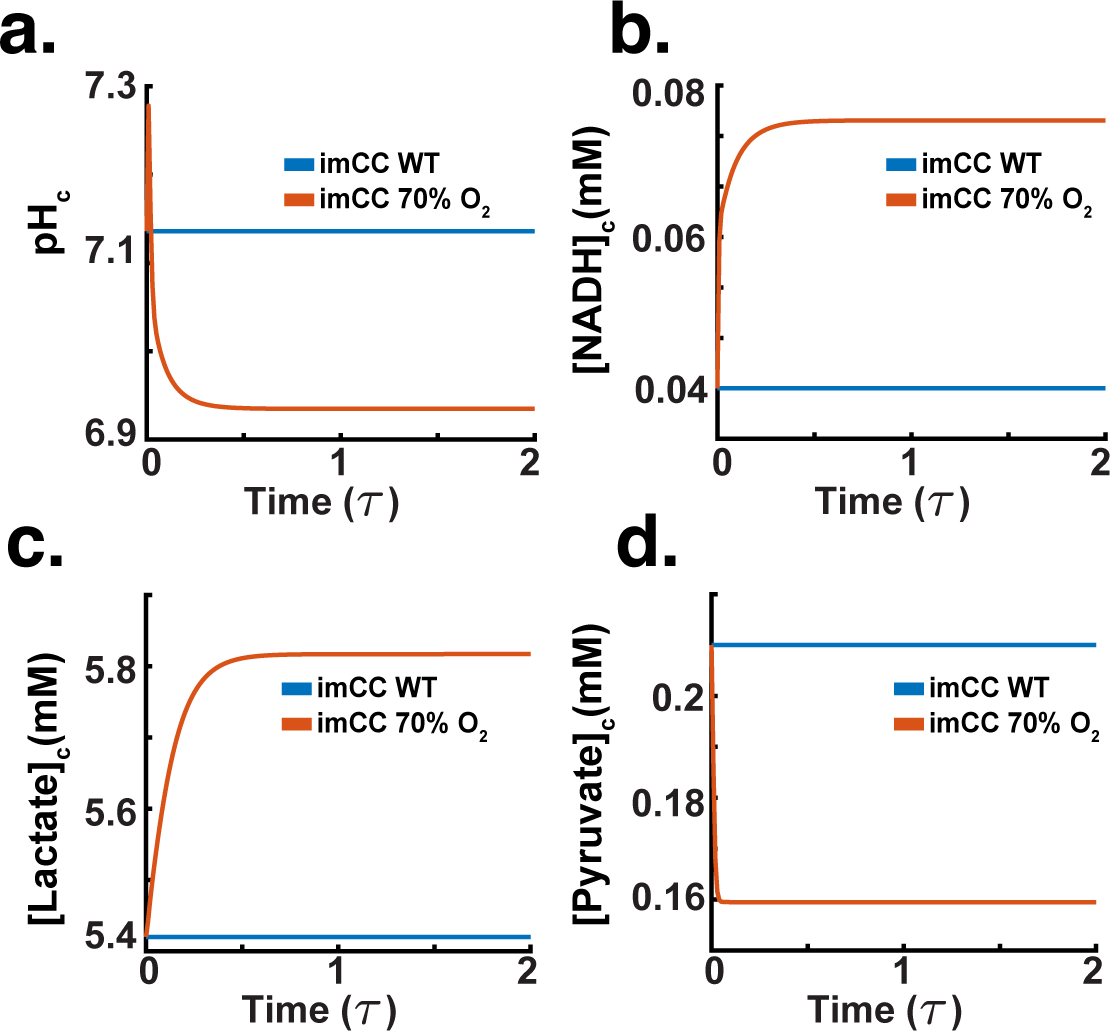
Benchmarking simulation (dimensonless time *τ*). We subjected the chromaffin cell model to hypoxia (a 30% reduction in oxygen flux). a. Acidification of the cytoplasm as a result of a metabolic shift towards anaerobic glycolysis. b. The cell exhibits an increase in NADH concentration, due to the inability of the glycolyytic pathway to rid lactate at the rate of normoxic respiration. c. Increase in lactate concentration due to a shift to anaerobic glycolysis. d. Decrease in pyruvate concentration is expected, most pyruvate is catabolised to generate lactate and thus mantain the cellular cofactor redox balance.

Based on these results (Figs. 10 and 11), the model has demonstrated a robust ability to replicate well-known metabolic shifts associated with mild hypoxia [78, 77, 1, 37, 38]. The observed changes in electron transport chain activity, lactate dehydrogenase activity, glycolysis, and intracellular pH align closely with documented physiological responses to hypoxic conditions [79, 78, 80].

### 5.2. SDH-b knockout

Given the validated performance of our model, we proceeded with simulations involving SDH-b knockouts to investigate the resulting metabolic consequences in chromaffin cells. Our model simulations reveal a comprehensive picture of the metabolic consequences of SDH-b knockout in chromaffin cells, particularly focusing on the activity of electron transport chain (ETC) complexes.

As shown in Fig. 12a, our model demonstrates that upon SDH-b loss, Complex I activity is laregly maintained, suggesting a compensatory mechanism wherein the cell has the capacity to retain Complex I activity to maintain electron flow into the ETC despite the impaired function of Complex II. This finding aligns with previous studies indicating cellular adaptation strategies to sustain mitochondrial function under compromised conditions [7, 11, 8, 23, 10]. The activities of Complex III and IV exhibited substantial decreases, with Complex III activity dropping by 60% and Complex IV activity by approximately 70% (Figs. 12b and 12c). These reductions indicate that the loss of electron supply from ubiquinol, due to the absence of SDH-b function, severely impacts downstream complexes. Despite the compensatory upregulation of Complex I, it is insufficient to fully sustain the activities of Complex III and IV. This observation is consistent with the known interdependencies within the ETC, where disruption in one complex can significantly affect the overall electron transport efficiency [82].

**Figure 12:**
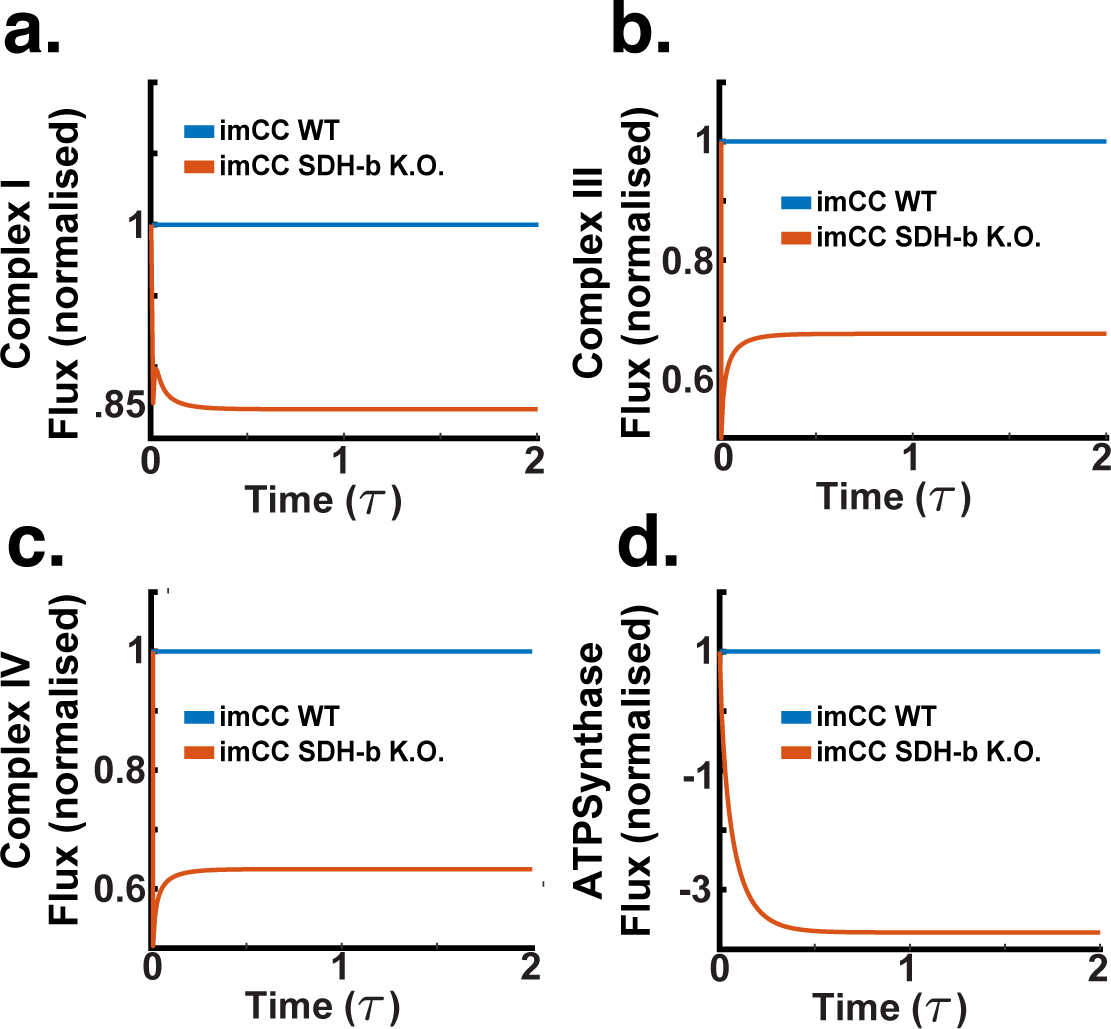
SDH-b knockout simulations (dimensionless time *τ*). a. Complex I activity is maintained after SDH-b loss, suggesting a compensatory upregulation to sustain electron flow into the ETC despite Complex II impairment. b.Complex III activity decreases by approximately 40%, indicating reduced electron supply from ubiquinol due to the loss of SDH-b function. c. Complex IV activity decreases by around 40%, further highlighting the impact on downstream complexes from the absence of SDH-b function. d. ATP synthase activity reverses from 1 to -4, showing a severe stress response where ATP synthase hydrolyses cytoplasmic ATP to maintain mitochondrial membrane potential, depleting cellular ATP reserves.

A significant observation was the reversal of ATP synthase activity, which shifted from 1 to -4 (Fig. 12d), as observed by [11]. This severe metabolic stress response indicates that ATP synthase hydrolyses cytoplasmic ATP to pump protons back into the mitochondrial matrix, a critical action to maintain the compartment’s membrane potential in the face of compromised ETC function [11]. This reversal comes at the cost of depleting cellular ATP reserves, highlighting the metabolic strain imposed by SDH-b knockout. To sustain this proton flux, the model relies heavily on proton leak mechanisms, which balance the activities of Complexes I through IV, reflecting the cell’s attempt to preserve mitochondrial integrity under stress.

Upon investigating cytosolic metabolite results, depicted in Fig. 13, we were able to further elucidate the metabolic consequences of SDH-b knockout in chromaffin cells. These observations highlight the pseudohypoxic phenotype characteristic of phaeochromocytomas, as reported by Kl’učková et al. [11].

**Figure 13:**
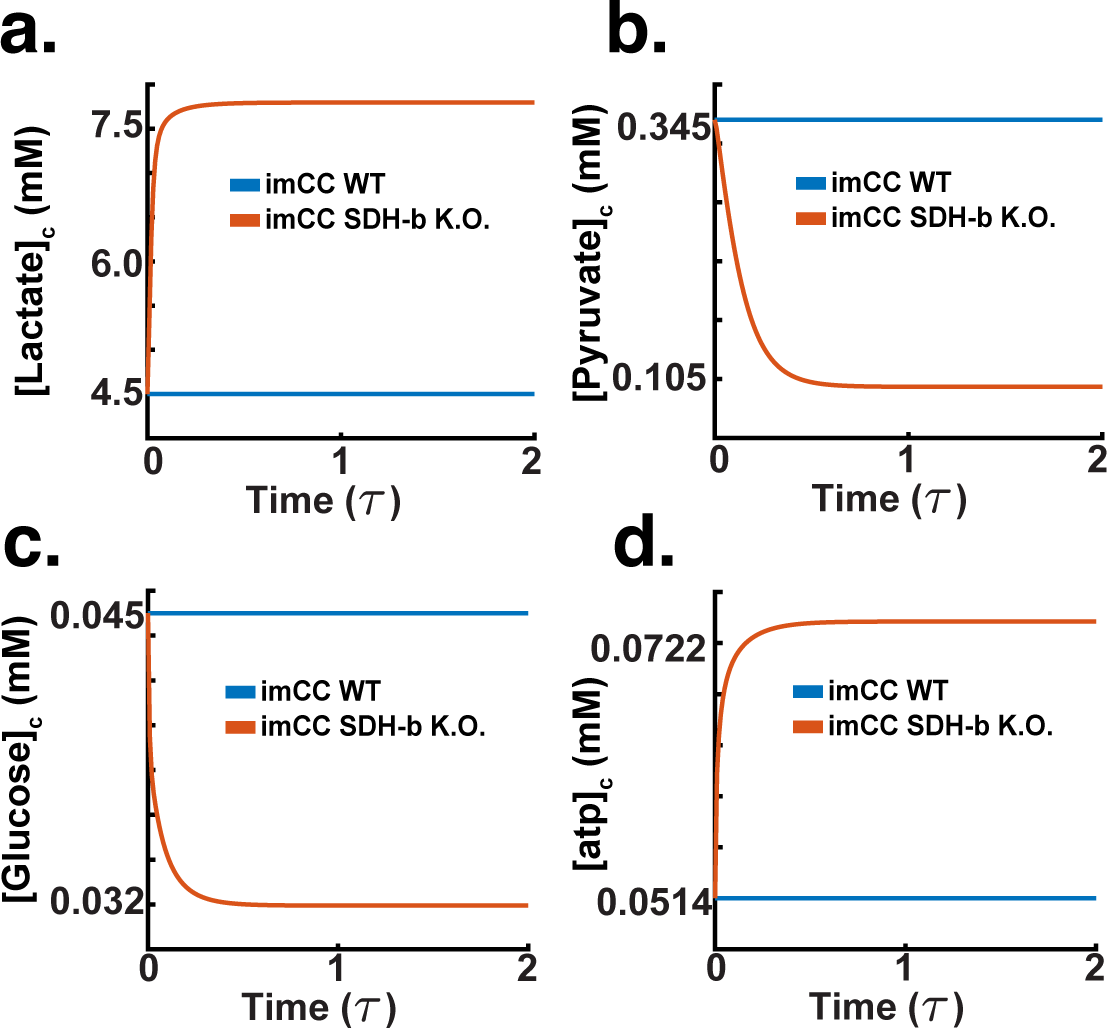
SDH-b knockout simulations (dimensionless time *τ*). a. Intracellular lactate concentration increases in SDH-b knockout cells, indicating a shift towards anaerobic glycolysis despite unchanged oxygen flux. b. Intracellular pyruvate concentration decreases, suggesting accelerated conversion to lactate, supporting enhanced glycolytic flux. c.Intracellular glucose concentration decreases, indicating increased glucose uptake and utilisation through glycolysis, aligning with the Warburg effect observed in cancer cells. d. Intracellular ATP concentration maintenance, reflecting impaired ATP synthesis due to dysfunctional ETC activity and consistent with the observed ATP synthase reversal, exacerbating cellular energy stress.

Fig.13a shows a marked increase in intracellular lactate concentration in SDH-b knockout cells compared to wild-type (WT) cells. This substantial rise is indicative of a shift towards anaerobic glycolysis, as seen in our hypoxia benchmarking results. Although this is a common response to hypoxia, where cells rely more heavily on glycolysis for ATP production due to compromised oxidative phosphorylation, here the net respiration of the system is not reduced. In fact, the model’s elevated lactate levels corroborate the increased activity of lactate dehydrogenase observed in previous simulations (Fig.12b), reinforcing the model’s accuracy in mimicking hypoxic-like metabolic responses.

In Fig. 13b, it is possible to observe a significant reduction in intracellular pyruvate concentration in SDH-b knockout cells. This decrease suggests an accelerated conversion of pyruvate to lactate, further supporting the enhanced glycolytic flux. The reduced pyruvate levels align with the expected metabolic adjustments where pyruvate is rapidly utilised to maintain glycolytic ATP production under conditions where the ETC function is impaired [68].

Fig.13c illustrates a decrease in intracellular glucose concentration in SDH-b knockout cells. Looking at the flux rate, this result indicates an uptick in glucose uptake and utilisation through glycolysis, a compensatory mechanism to counteract the diminished ATP production from oxidative phosphorylation. This model also shows a decreased reliance on extracellular glucose, which aligns with our experimental data (Fig.5b) [11, 9, 78].

Lastly, Fig.13d shows a maintenance of intracellular ATP concentration in SDH-b knockout cells relative to WT, reflecting the hydrolysis of mitochondrial ATP due to dysfunctional ETC activity this occurs as a compensatory mechanism via the glycolytic pathway. The ATP levels are consistent with the observed reversal of ATP synthase activity by Kl’učková et al. [11] (Fig.12d), where they observed ATP hydrolysis occurs to maintain mitochondrial membrane potential via blockage of ATPsynthase using olygomycin.

The observed shift to a more glycolytic phenotype despite normoxic conditions has important consequences in mitochondrial metabolism. In Fig.14a, the model indicates that the concentration of mitochondrial pyruvate upon SDH-b knockout is significantly reduced compared to WT cells. This reduction aligns with the earlier observed decrease in cytosolic pyruvate (Fig.13b) and suggests a disruption in pyruvate import into the mitochondria, likely due to impaired mitochondrial function and a shift towards cytosolic metabolism [79, 80].

Fig.14b shows the concentration of mitochondrial succinate. In SDHb knockout cells, succinate levels are markedly elevated compared to WT cells. This accumulation of succinate is a hallmark of SDH deficiency and is consistent with previous findings that link elevated succinate levels to the stabilisation of hypoxia-inducible factors (HIFs), promoting a pseudohypoxic phenotype even under normoxic conditions [11, 7, 13, 10, 8]. Note that the model indicates that the elevated succinate levels impede the proper functioning of the electron transport chain, further corroborating the observed reductions in Complex III and IV activities (Figs.12b and 12c).

The model also shows a decrease in fumarate concentration, as seen in Fig. 14c. The decrease in fumarate is an expected consequence due to the bottleneck created by the loss of the SDH-b subunit, which compromises succinate dehydrogenase (SDH) activity. However, there is a slight accumulation of downstream metabolites like malate and fumarate. The buildup of these intermediates results from pyruvate carboxylases redirecting the flux of carbons towards the malate/aspartate shuttle. The model indicates this supports the increased oxidation of NADH into NAD^+^. This disrupts the TCA cycle and contributes to metabolic shifts associated with the pseudohypoxic response observed in chromaffin cells upon SDH-b loss [11].

**Figure 14:**
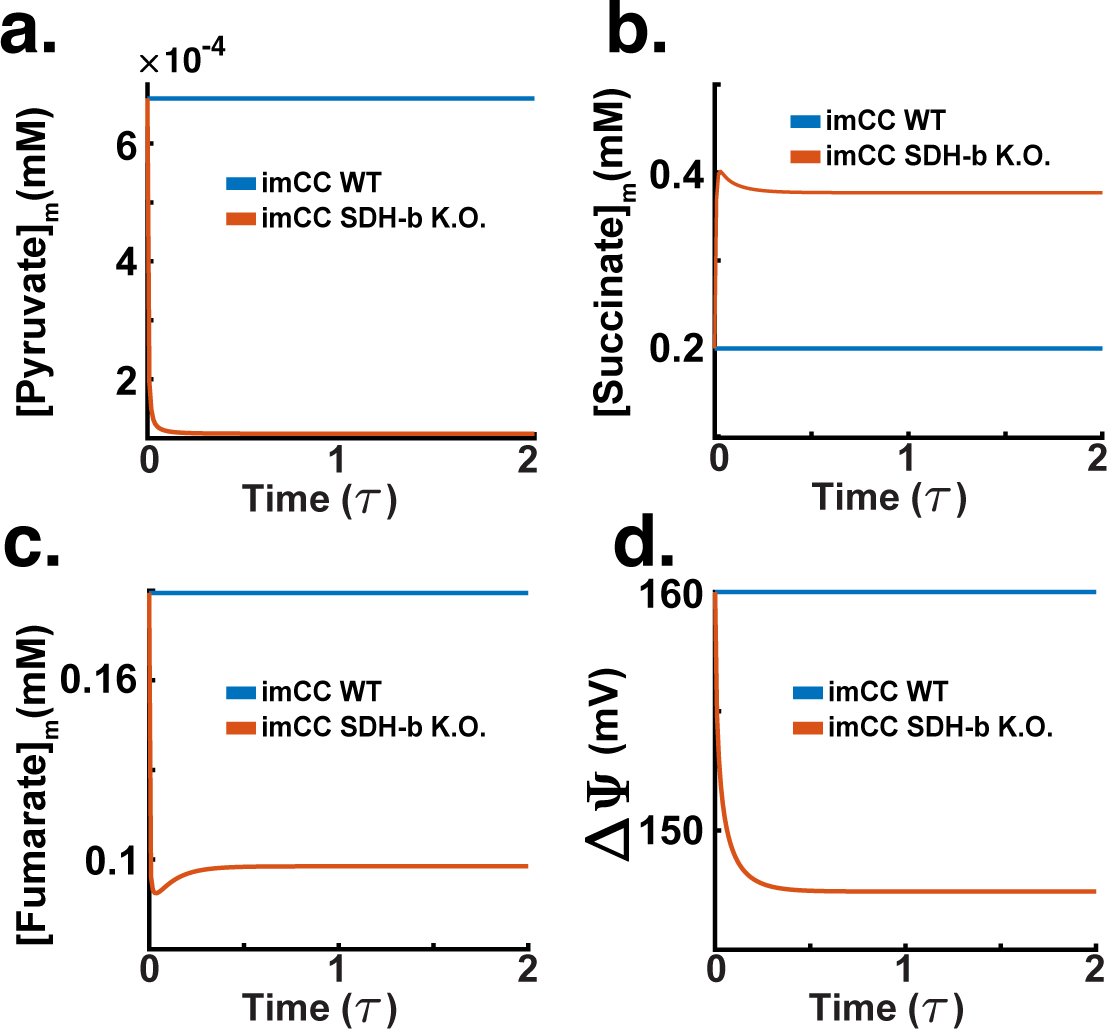
SDH-b knockout simulations (dimensionless time *τ*). a. Mitochondrial pyruvate concentration significantly decreases in SDH-b knockout cells, indicating disrupted pyruvate import and a shift towards cytosolic metabolism. b. mitochondrial succinate concentration increases markedly in SDH-b knockout cells, a hallmark of SDH deficiency, promoting a pseudohypoxic phenotype by stabilising hypoxia-inducible factors. c. Mitochondrial fumarate concentration decreases due to the loss of SDH-b, creating a bottleneck in the TCA cycle, with slight upstream accumulation of intermediates like malate and fumarate. d.Mitochondrial membrane potential (ΔΨ) decreases, reflecting impaired electron transport chain function and energy stress.

The mitochondrial membrane potential (ΔΨ) significantly decreases in SDH-b knockout cells, as shown in Fig. 14d. This reduction in membrane potential indicates impaired electron transport chain (ETC) function. The diminished (ΔΨ) reflects the compromised ability of the ETC to maintain proton gradients across the mitochondrial membrane, crucial for ATP synthesis. This loss of potential, the model indicates, is a direct consequence of the disrupted function of Complex II. The impaired ETC function exacerbates cellular energy stress, forcing the cell to rely more heavily on anaerobic glycolysis to meet its energy demands. The decrease in (ΔΨ) further supports the observation of reversed ATP synthase activity (Fig. 12d), where ATP is hydrolysed to pump protons back into the mitochondrial matrix, a desperate measure to preserve mitochondrial integrity and function under stress.

### 5.3. Mitochondrial swelling

In addition to the metabolic changes highlighted by our simulations, the model also highlights significant alterations in mitochondrial volume and the proton motive force (PMF) in SDH-b knockout cells.

The results above do not reflect the full consequence of a reversal of the ATP synthase. The model indicates that this activity has significant implications, evident upon examining the model’s proton motive force (PMF) across the mitochondrial membrane. Briefly, the PMF is the electrochemical gradient generated by the ETC, consisting of two components: a pH gradient (difference in proton concentration) and an electrical potential gradient (voltage difference) across the inner mitochondrial membrane [83, 19]. In normal conditions, ATP synthase utilises the PMF to drive the synthesis of ATP from ADP and inorganic phosphate. Protons flow back into the mitochondrial matrix through ATP synthase, which drives the phosphorylation of ADP. When ATP synthase operates in reverse (due to severe metabolic stress or impaired ETC function), it hydrolyses ATP to pump protons from the mitochondrial matrix to the intermembrane space. This reversal is a compensatory mechanism to maintain the PMF, particularly the electrical potential gradient [18].

The reversal of ATP synthase activity, in this model, helps to maintain the electrical component of the PMF. By hydrolysing ATP and pumping protons out of the matrix, ATP synthase helps sustain the voltage difference across the inner mitochondrial membrane. However, this comes at the cost of depleting the available mitochoindrial ATP pool, as ATP is consumed to sustain the proton gradient instead of being produced. The pH gradient (difference in proton concentration) is also affected. Proton pumping by reversed ATP synthase increases the proton concentration in the intermembrane space, which can help maintain the pH gradient [18]. Despite this, the overall efficiency of the proton gradient might be reduced due to the impaired ETC function, which normally contributes significantly to proton pumping. The combined effect of ATP synthase reversal and reduced ETC efficiency results in a complex balance. The PMF is partially maintained by the reversed ATP synthase activity, but the system is under stress, leading to an overall less stable and energetically costly maintenance of the PMF.

Fig. 15a shows the model’s reaction to SDH-b loss by demarcating a decrease in mitochondrial NADH concentration compared to WT imCCs. This decrease indicates impaired oxidative phosphorylation, as NADH is a crucial substrate for the ETC function, and its reduced levels reflect, in this model, a diminished electron transfer through the chain as seen in our results above (Figs. 13 and 14). However, the result is also indicative of a bottleneck in the TCA cycle, as the uncoupling of the malate/aspartate shuttle prevents the model from oxidising NAD^+^ for its use in the electron transport chain. These results reflect the metabolic interplay seen in our own experimental data (Fig. 7)

**Figure 15:**
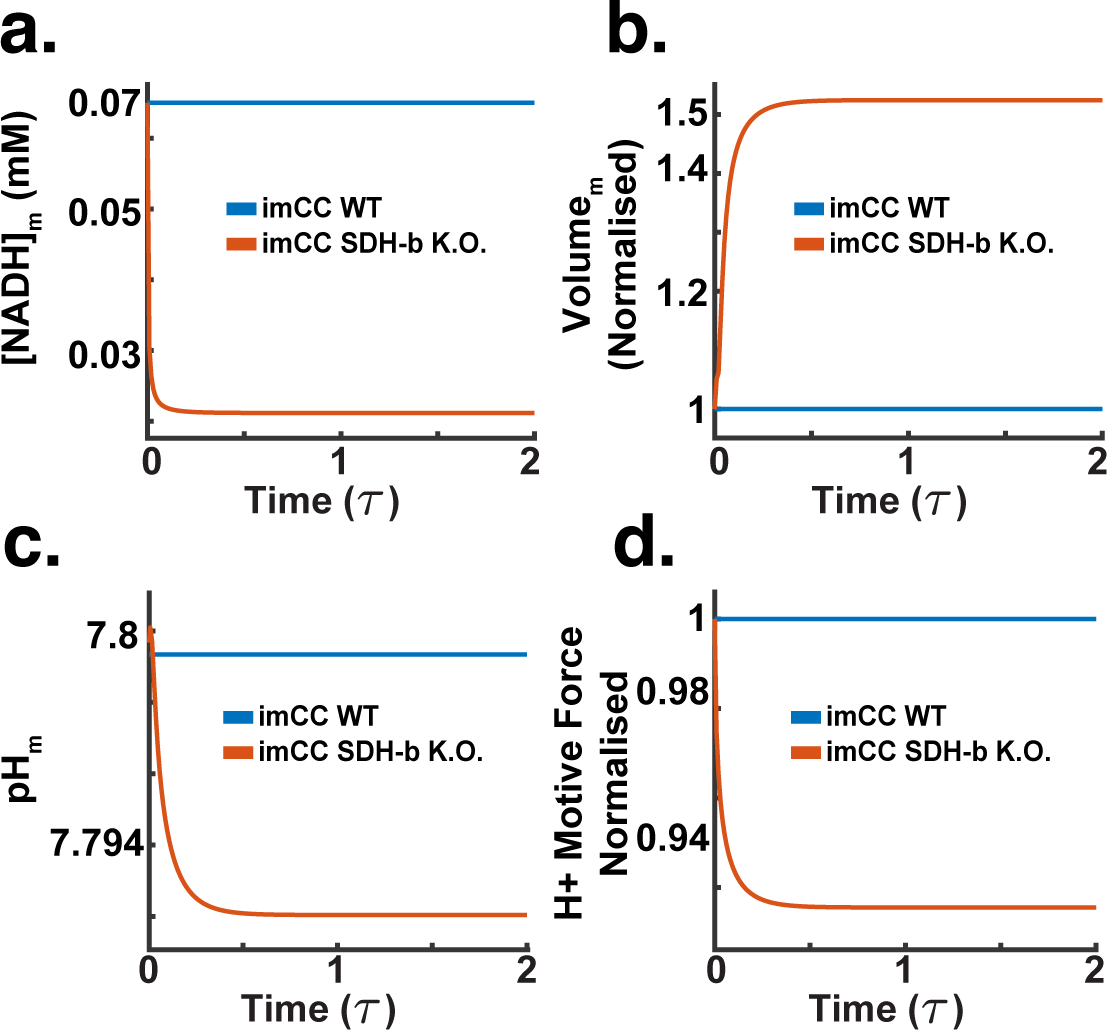
SDH-b knockout simulations (dimensionless time *τ*). a. Mitochondrial NADH concentration decreases in SDH-b knockout cells, indicating impaired oxidative phosphorylation and a bottleneck in the TCA cycle. b.Mitochondrial volume increases in SDH-b knockout cells, reflecting altered osmotic balance and ionic homeostasis due to disrupted ETC function.c. Mitochondrial pH slightly decreases, consistent with increased proton concentration in the intermembrane space caused by reversed ATP synthase activity and impaired ETC function.d. Proton motive force (PMF) decreases in SDH-b knockout cells. Despite compensatory reversal of ATP synthase activity, the PMF is maintained but not at optimal levels, indicating energetically costly maintenance of the electrochemical gradient.

In Fig. 15b, it is possible to see how the loss of SDH-b leads to an increase in mitochondrial volume. This increase, in our model, is a direct response to the altered osmotic balance and ionic homeostasis resulting from disrupted ETC function. Mitochondrial swelling is a well-documented response to metabolic stress caused by SDH-b loss [11, 8]. Unlike observations by Kl’učková et al. [11], our model did not achieve the expected fold-change. This could be due to diverse factors, we suspect our model does not account for other metabolic alterations that may affect mitochondrial swelling (i.e., the mitochondrial transition pore mPTP, which is regulated by Ca^2+^).

Concomitantly, the model shows a slight decrease in mitochondrial pH in SDH-b knockout cells (Fig. 15c). The decrease in pH is consistent with the increased proton concentration in the mitochondrial intermembrane space due to the reversed ATP synthase activity and impaired ETC function. However, this acidification is not too drastic, as might be naively expected. This is due, the model indicates, to its capacity of sustaining the proton motive force under metabolic stress (i.e., ATP synthase reversal).

Indeed, Fig. 15d depicts a decrease in the proton motive force (PMF) in SDH-b knockout cells. Despite the compensatory reversal of ATP synthase activity, the PMF is maintained, albeit not at optimal levels. The reduced PMF indicates an overall less efficient and energetically costly maintenance of the electrochemical gradient, crucial for ATP production and mitochondrial function.

### 5.4. Complex 1 retention

As the SDH-b’s ability to transport electrons through its sulfur-iron clusters (denoted in the model as *k*_[_3*_Fe−_*4*_S_*_]_) decreases, initially, there is an observed increase in Complex I (C1) activity. This increase in C1 activity occurs as a compensatory response by the cell to maintain electron flow into the ETC, even as the function of Complex II is impaired due to SDH-b loss this observation is in accordance to what Kl’učková et al. [11] report. Furthermore, this adaptive mechanism, the model reveals, helps in sustaining the proton motive force (PMF), ensuring that ATP synthesis can continue, albeit at a potentially reduced efficiency [18].

However, as the capacity of SDH-b continues to decline beyond a critical point, the compensatory mechanism of upregulating Complex I becomes insufficient. This is depicted in Fig. 16 where the PMF begins to plateau and eventually declines as the ability of the enzyme to transport electrions (all the way to SDH-b K.O.) diminishes. As a result, the cell can no longer sustain the necessary proton gradient solely through Complex I activity, leading to a total collapse of the ETC functionality.

**Figure 16:**
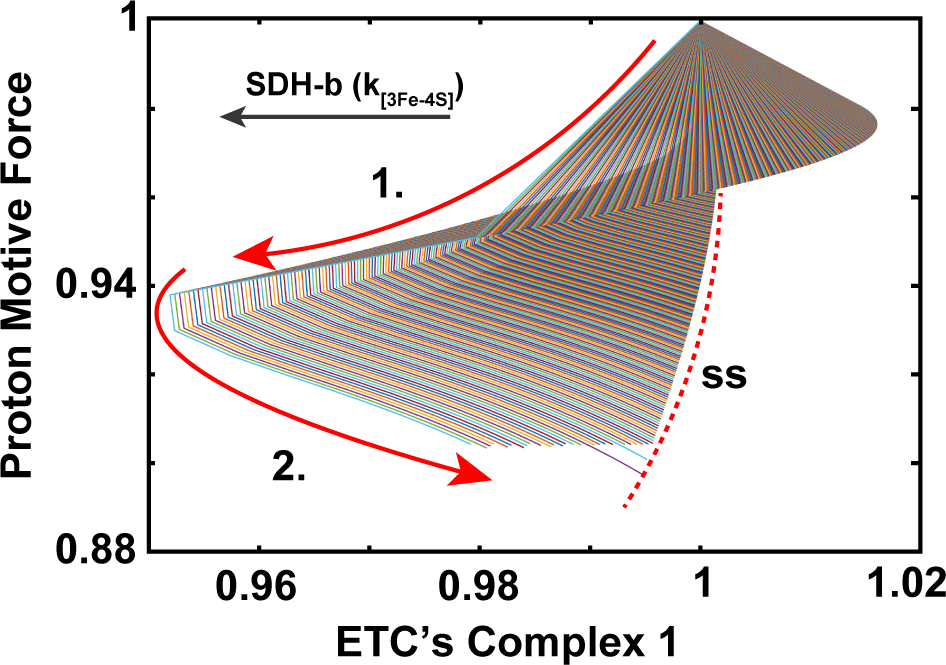
SDH-b knockout simulation. a. Surface plot showing the relationship between Proton Motive Force (PMF) and ETC Complex I activity as SDH-b electron transport capacity (*k*_3*F*_ *_e−_*_4*S*_) decreases. Initially, Complex I activity increases as a compensatory mechanism to sustain PMF.

This result highlights the initial adaptive increase in Complex I activity in response to SDH-b loss, followed by a collapse in the system as the compensatory mechanisms are overwhelmed. The maintenance of PMF through proton leaks and the reversal of ATP synthase activity underscores the cell’s efforts to adapt to metabolic stress and sustain mitochondrial function under compromised conditions.

Our previous study (Kl’učková et al. [11]) reported that despite SDHb deficiency, chromaffin cells retain Complex I function, which is critical for maintaining metabolic fitness. Their study shows that the retention of Complex I activity helps to sustain mitochondrial NADH oxidoreductase function, thereby preventing a complete metabolic collapse. This supports our model’s prediction of an initial compensatory upregulation of Complex I activity, followed by a critical threshold beyond which the system fails to maintain ETC functionality.

In-depth discussion of the surface plot reveals key insights into the cellular adaptations and limitations in response to SDH-b deficiency. Initially, as SDH-b function decreases, the electron transport chain compensates by upregulating Complex I activity to maintain the PMF. This adaptive response is reflected in the transient increase in Complex I activity, which helps to sustain ATP production and cellular energy balance. However, this compensatory mechanism has its limits. As SDH-b capacity further declines, the system reaches a tipping point where Complex I alone cannot sustain the electron flow and PMF, leading to a gradual decline in ETC efficiency.

Fig. 16 highlights the dynamic interplay between SDH-b activity and Complex I function. The observed fold in the solution surface indicates a critical threshold beyond which the compensatory mechanisms can no longer cope with the metabolic stress. At this point, the system experiences a rapid decline in ETC function, leading to impaired mitochondrial respiration and energy production all of this whilst the proton leak salvages the PMF. This fold represents a ‘sink’ in the mathematical sense, where the system’s state is drawn towards a stable, but suboptimal, condition of impaired Complex I activity and reduced PMF. This proton leaks seemingly, is able to dissipate the proton gradient, reducing the efficiency of ATP synthesis but preventing a complete collapse of the mitochondrial membrane potential. This leak mechanism, along with the reversal of ATP synthase activity, highlights the cell’s efforts to adapt to the severe metabolic stress induced by SDH-b loss. The reliance on proton leaks and ATP synthase reversal indicates that this compensation comes at a significant energetic cost, the model predicts this could limit the cells’ ability to cope with additional stressors despite their perceived metabolic robustness.

## 6. Discussion

The primary objective of this study was to develop a comprehensive mathematical model to elucidate the metabolic consequences of SDH-b loss in chromaffin cells, particularly focusing on the electron transport chain (ETC) and associated metabolic pathways. Our findings, derived from detailed simulations and experimental data integration, provide significant insights into the adaptive mechanisms employed by chromaffin cells in response to SDH-b deficiency [11].

To validate and extend previous observations, we revisited ^13^C_6_-glucose labelling experiments, identifying any discrepancies or inconsistencies between our current and past findings. This approach facilitated a deeper understanding of the underlying biological processes driving our mathematical model (Fig. 6 and 7). By subjecting the cells to fully labelled glucose (^13^C_6_-glucose), we aimed to gather comprehensive data on metabolite concentrations, flux rates, and enzyme kinetics (Fig. 8 to 10). These data serve as crucial inputs for refining model parameters, ensuring their congruence with experimental observations.

Our model simulations support earlier observations that SDH-b knockout leads to notable alterations in the activity of ETC complexes [11, 8, 10]. While Complex I activity is maintained through compensatory upregulation, Complexes III and IV exhibit substantial decreases in activity (Fig. 13). This imbalance underscores the critical interdependencies within the ETC, where impairment in one complex can propagate through the chain, affecting overall electron transport efficiency. The maintenance of Complex I activity suggests that cells may activate alternative pathways or regulatory mechanisms to uphold mitochondrial function under stress conditions, aligning with previous studies on cellular adaptation strategies [7, 11].

A significant finding in our simulations was the reversal of ATP synthase activity, where the enzyme hydrolyses ATP to pump protons back into the mitochondrial matrix (Fig. 12d). This mechanism, the model indicates, serves to maintain the mitochondrial membrane potential in the face of compromised ETC function but at the cost of cellular ATP reserves (Fig. 14d). This reversal is indicative of severe metabolic stress and highlights the cell’s prioritization of maintaining membrane potential over efficient ATP production [11].

SDH-b knockout simulations also elucidated a shift towards a pseudohypoxic phenotype, characterised by increased lactate production and reduced pyruvate and glucose levels (Fig. 13). These changes reflect an enhanced reliance on glycolysis for ATP production, mimicking the Warburg effect commonly observed in cancer cells [8, 78]. The elevated intracellular lactate and reduced pyruvate concentrations align with a metabolic shift towards anaerobic glycolysis, even under normoxic conditions. This shift is a well-documented response to hypoxia, but in our model, it occurs as a direct consequence of SDH-b loss, independent of oxygen availability [7, 8, 10].

Our results also observed a reproduction of experimental data where upon SDH-b knockout, an increase in mitochondrial volume led to irreversible changes in the cell’s proton motive force (PMF) (Fig. 15). The model indicated that mitochondrial swelling occurs as a response to altered osmotic balance and ionic homeostasis resulting from disrupted ETC function. The maintenance of PMF, despite the reversal of ATP synthase activity, underscores the cell’s effort to preserve mitochondrial integrity under stress. However, this comes at the cost of increased energetic demands, highlighting the metabolic strain imposed by SDH-b knockout [11, 8].

The mathematical model presented in this study provides a first-step towards the construction of a robust framework for understanding the metabolic consequences of SDH-b loss in chromaffin cells. The findings highlight the cell’s adaptive mechanisms, including the compensatory upregulation of Complex I, the reversal of ATP synthase activity, and the metabolic shift towards a pseudohypoxic phenotype, a poorly understood phenomena seen in phaeochromocytomas. These insights not only advance our understanding of the metabolic adaptations in SDH-b deficient cells but also offer potential avenues for new insights in other conditions where SDH plays a centre role. In subsequent studies we aim to refine the model by incorporating additional regulatory mechanisms, like incorporating the intersection of cell-signalling and metabolism (Ca^2+^) and validating the findings with further experimental data.

## 7. CRediT author statement

**Eĺıas Vera-Sigüenza:** Mathematical Model Conceptualisation, Computational Methodology, Software,Visualisation, Model Validation, Model Analysis, Writing, Editing. **Himani Rana:** Experimental Data Procurement. **Ramin Nashebi:** Visualisation, Model Validation, Model Analysis **Ielyaas Cloete:** Visualisation, Model Validation, Model Analysis **Kataŕına Kl’učková:** Conceptualisation, Experimental Data Procurement. **Fabian Spill:** Article Review & Editing. **Daniel A. Tennant:** Study Conceptualisation, Experimental Data Validation, Article Editing, Funding acquisition.

## Supporting information

Experimental Data (labelling experiments)

## Acknowledgements

ES, HR, JF, and DA.T are supported by the Paradifference Foundation. IC is supported by a Maŕıa de Maeztu Post-doctoral Fellowship. ES, HR, and DA.T are supported also by a Cancer Research UK Grant (C42109/ A26982 and C42109/A24891). FS and RN are supported by a UKRI Future Leaders Fellowship Grant (MR/T043571/1)

