## Supplement Material for "A Mathematical Exploration of SDH-b Loss in Chromaffin Cells"

**Abstract**

The following are brief descriptions of the formulation of the equations of the model.

### 1. Nkcc Co-transporter

The Nkcc co-transporter is ubiquitously expressed in nearly all cells [1]. The Nkcc-mediated $\mathrm{Cl}^-$ uptake mechanism is a secondary active transport process, meaning the energy required for its activity is indirectly derived from ATP hydrolysis. The Nkcc model we use was initially constructed by Benjamin and Johnson [2]. This model assumes equilibrium ion binding, binding symmetry, and identical $\mathrm{Cl}^-$ binding sites, resulting in a ten-state model. Palk et al. [3],Vera-Sigüenza et al. [4] and Gin et al. [5] simplified it to a two-state model that assumes simultaneous binding and unbinding of $\mathrm{Cl}^-$, $\mathrm{Na}^+$ and $\mathrm{K}^+$. The reaction proceeds as follows:

$$O + \mathrm{K}_e^+\mathrm{Na}_e^+(\mathrm{Cl}_e)^2 \underset{k_1^-}{\overset{k_1^+}{\rightleftharpoons}} I \underset{k_2^-}{\overset{k_2^+}{\rightleftharpoons}} O + \mathrm{K}_i^+\mathrm{Na}_i^+(\mathrm{Cl}_i^-)^2.$$

The steady-state flux is given by:

$$J_{\mathrm{Nkcc1}} = \alpha_{\mathrm{Nkcc1}} \left( \frac{a_1 - a_2[\mathrm{Na}^+]_i[\mathrm{K}^+]_i[\mathrm{Cl}^-]_i^2}{a_3 + a_4[\mathrm{Na}^+]_i[\mathrm{K}^+]_i[\mathrm{Cl}^-]_i^2} \right), \tag{1}$$

where $\alpha$Nkcc1 is the density of the co-transporter.

### 2. Chloride and Potassium Channels

Previous mathematical models have used linear relationships in the uptake and extrusion of $Cl^-$. We adapted the model of Arreola et al. [6]. Although we found no major qualitative difference between models [4], our choice of model, however, predicts a large maximum single channel conductance as a function of membrane potential. The chloride flux is defined as

$$J_{\mathrm{Cl}} = \frac{G_{\mathrm{Cl}}}{F}\left(V_m - V_{\mathrm{Cl}}\right), \tag{2}$$

where the Nernst potential is given by:

$$V_{\mathrm{Cl}} = \frac{RT}{z^{\mathrm{Cl}}F}\ln\left(\frac{[\mathrm{Cl}^-]_e}{[\mathrm{Cl}^-]_c}\right).$$

Here $z^{Cl} = -1$, the ion's valence. In a similar vein, we modelled the membrane potassium fluxes as: s

$$J_{\mathrm{K}} = \frac{G_{\mathrm{K}}}{F}\left(V_m - V_{\mathrm{K}}\right), \tag{3}$$

with Nernst potential

$$V_{\mathrm{K}} = \frac{RT}{Fz^{\mathrm{K}}}\ln\left(\frac{[\mathrm{K}^+]_e}{[\mathrm{K}^+]_c}\right),$$

where $z^{\mathrm{K}} = +1$, the ion's valence.

### 3. NaK ATPase Pump

The NaK ATPase pump extrudes 3 $Na^+$ ions while importing 2 $K^+$ ions against their electrochemical gradients, utilising one ATP molecule per cycle. The net reaction for the pump cycle is:

$$\mathrm{ATP} + 3\mathrm{Na}_i^+ + 2\mathrm{K}_e^+ \leftrightarrow \mathrm{ADP} + \mathrm{P}_i + 3\mathrm{Na}_e^+ + \mathrm{K}_i^+,$$

where $P_i$ denotes the intracellular concentration of phosphate, a byproduct of ATP conversion to ADP. Crampin et al. [7], Smith and Crampin [8] developed a mathematical model of the NaK-ATPase pump intended for use in whole-cell myocyte modelling, with the goal of predicting pump function and overall cell behaviour when cellular metabolism is impaired. Palk et al. [3] simplified this model to a two-state system:

$$\mathrm{O} + 2\mathrm{K}_e + \mathrm{ADP}_e + \mathrm{P}_{\mathrm{i}e} \underset{k_1^-}{\overset{k_1^+}{\rightleftharpoons}} \mathrm{I} + 3\mathrm{Na}_e + \mathrm{ATP}_e,$$

$$\mathrm{I} + 3\mathrm{Na}_i + \mathrm{ATP}_c \underset{k_2^-}{\overset{k_2^+}{\rightleftharpoons}} \mathrm{O} + \mathrm{ADP}_c + \mathrm{P}_{\mathrm{i}c} + 2\mathrm{K}_i.$$

Here 'I' refers to the 'Inside' state and 'O' to the 'Outside' state. This simplification assumes the simultaneous binding and unbinding of external $Na^+$ and internal $K^+$ ions, supplied at a constant rate. It also assumes that the forward reaction rates exceed the reverse rates. The steady-state flux through the pump is expressed as:

$$J_{\mathrm{NaK}} = \alpha_{\mathrm{NaK}} \left( \frac{[\mathrm{ADP}]_{\mathrm{e}}[\mathrm{K}^+]_{\mathrm{e}}^2[\mathrm{Na}^+]_{\mathrm{c}}^3[\mathrm{ATP}]_{\mathrm{c}}}{[\mathrm{ADP}]_{\mathrm{e}}[\mathrm{K}^+]_{\mathrm{e}}^2 + \alpha_{\mathrm{NaK}}[\mathrm{Na}^+]_{\mathrm{c}}^3[\mathrm{ATP}]_{\mathrm{c}}} \right), \tag{4}$$

where $\alpha_{\mathrm{NaK}}$ is the density of the pump [9].

### 4. Carbonic Anhydrases

The cell's membrane is permeable to $CO_2$ which diffuses down its concentration gradient into the cytoplasm where it combines with water to form carbonic acid ($H_2CO_3$). Carbonic anhydrases catalyse the reaction and dissociate the acid quickly into $H^+$ and $HCO_3^-$. Here we assume the reaction occurs as follows:

$$\mathrm{CO_2} + \mathrm{H_2O} \underset{k_{-1}}{\overset{k_1}{\rightleftharpoons}} \mathrm{H}^+ + \mathrm{HCO_3^-}.$$

Using the law of mass action we derived a term for the production of $HCO_3^-$ and $H^+$ that is proportionally dependent on the concentration of $CO_2$,

$$J_{\mathrm{Buffer}} = k_1[\mathrm{CO_2}]_c - k_{-1}[\mathrm{H}^+]_c[\mathrm{HCO_3^-}]_c. \tag{5}$$

### 5. Na/H exchanger (Nhe)

We used a model developed by Falkenberg and Jakobsson [10], which utilises the $HCO_3^-$ concentration gradient to drive the influx of $Cl^-$ into the cell. The energy for this ion exchange is provided by NaK-ATPase activity [11]. The flux of this model is influenced by the ionic concentrations in the cytoplasm and interstitial fluid, the number of available binding sites, and the half-saturation constants. The model calculates the flux as the product of two factors: the ionic concentration contributions and the binding site characteristics, described by Michaelis-Menten kinetics, and the conductance of the exchanger, which is proportional to the density of active membrane proteins. For simplification, we incorporate the reverse operation of the transporter, making the exchanger bidirectional:

$$J_{\mathrm{Nhe1}} = G_{\mathrm{Nhe1}} \Bigg[ \left( \frac{[\mathrm{H}^+]_c}{[\mathrm{H}^+]_c + K_{\mathrm{H}}} \right)^2 \left( \frac{[\mathrm{Na}^+]_e}{[\mathrm{Na}^+]_e + K_{\mathrm{Na}}} \right) - \left( \frac{[\mathrm{Na}^+]_c}{[\mathrm{Na}^+]_c + K_{\mathrm{Na}}} \right) \left( \frac{[\mathrm{H}^+]_e}{[\mathrm{H}^+]_e + K_{\mathrm{H}}} \right)^2 \Bigg]. \tag{6}$$

### 6. Chloride-Bicarbonate Antiporter (Ae2)

Similarly to the Nhe antiporter, our model for the Ae2 exchanger is based on Falkenberg and Jakobsson [10]. Its flux is given by

$$J_{\mathrm{Ae2}} = G_{\mathrm{Ae2}} \Bigg[ \left( \frac{[\mathrm{Cl}^-]_e}{[\mathrm{Cl}^-]_e + K_{\mathrm{Cl}}} \right) \left( \frac{[\mathrm{HCO}_3^-]_c}{[\mathrm{HCO}_3^-]_c + K_{\mathrm{B}}} \right) - \left( \frac{[\mathrm{Cl}^-]_c}{[\mathrm{Cl}^-]_c + K_{\mathrm{Cl}}} \right) \left( \frac{[\mathrm{HCO}_3^-]_e}{[\mathrm{HCO}_3^-]_e + K_{\mathrm{B}}} \right) \Bigg]. \tag{7}$$

### 7. Volume Model

It is a constraint of the model that the cellular compartments must maintain electroneutrality. The number of moles of large negatively charged molecules (with valence $z_x \leq -1$) that are impermeable to the cellular membrane and thus trapped inside the cell is denoted as $x_c$. To find its value, we note:

$$\sum_j [\text{ion}]_{c_j} - \frac{x_c}{\omega_c} = 0,$$

where $[\text{ion}]_{c_j}$ is the $j^{th}$ ion/metabolite, and its associated charge. $\omega_c$ is the volume of the cell. Solving for $x_c$:

$$x_c = \omega_c \sum_j [\text{ion}]_{c_j}.$$

This follows for all compartments of our cell.

Having obtained this value, we model compartmental volume as a direct consequence of the osmotic gradient between neighbouring compartments, assuming that the plasma membrane cannot withstand hydrostatic pressure gradients. The difference between the extracellular, mitochondrial, and cytoplasmic osmotic gradients leads to a change in cellular volume($\omega_i$):

$$\frac{d\omega_i}{dt} = q_a - q_b. \tag{8}$$

The terms $q_a$ and $q_b$ represent the osmotic pressure across the membranes separating these compartments, defined by:

$$q_a = P_a \left[ \sum [c]_e + \Phi_e - \sum [c]_c - \frac{x_c}{\omega_c} \right],$$

$$q_b = P_b \left[ \sum [c]_c + \frac{x_c}{\omega_c} - \sum [c]_m - \frac{x_m}{\omega_m} \right].$$

Here, $P_i$ ($a$ or $b$) represents the permeability of the respective plasma membrane portion, and the values $x_c$ the moles of negatively charged particles with valence $(z_x) \leq -1$ in the cellular media that are impermeable to the plasma membrane. To obtain its value, we used the electroneutrality condition described above. The term $\Phi_e$ accounts for the uncharged impermeable ionic species and proteins (e.g., amylase) present in the extracellular compartment that we do not keep track of in the model but contribute to the osmolarity of the compartment. This term is a parameter of the model and its value was found by solving the equation at steady state.

### 8. Membrane Potential

This model deals with different membrane potentials on different compartments of the cell; the cellullar membrane , the inner mitochondrial membrane (facing cytosol), and the mitochondrial matrix membrane - facing the inner mitochondrial space. To obtain the potential at each membrane portion we use Kirchhoff's current law for a simple resistor-capacitor circuit. For the cellular membrane we have:

$$C_m \frac{\mathrm{d}V_m}{\mathrm{d}t} = \sum I_{\mathrm{ion}}. \tag{9}$$

The right hand side represents the sum of the currents for each ion species through the respective membrane. $C_m$ is the capacitance of the cell membrane. We used the convention that the membrane potential is negative as it is measured from the extracellular space to the cytoplasm.

The model for the mitochondrial membrane potential in our model is directly taken from Beard [12]. Here the mitochondrial membrane potential ($\Delta\Psi$) is measured as the difference between inner mitochondrial membrane (facing cytosol) and the mitochondrial matrix membrane:

$$\Psi_m = -0.65\Delta\Psi,$$
$$\Psi_i = 0.35\Delta\Psi.$$

In this way we have that,

$$C_{m_i} \frac{\mathrm{d}\Delta\Psi}{\mathrm{d}t} = \sum I_{\mathrm{ion_m}}, \tag{10}$$

where $C_{m_i}$, is the effective membrane capacitance.

### 9. Gucose Transport (GLUT1)

To model the uptake of glucose, we opted for a Hill function type model [13, 14]. The flux or rate of glucose transport through the GLUT1 transporter is proportional to a scaling factor ($g_{GLUT}$) that adjusts the overall transport rate - the so called $V_{max}$. The term $\frac{K_{glc}[Glc]_e}{M_{glc}+[Glc]_e}$ models the Michaelis-Menten kinetics, where $[Glc]_e$ is the extracellular glucose concentration,$K_{glc}$is a constant related to glucose binding affinity, and$M_{glc}$is the Michaelis constant indicating the glucose concentration at which the transport rate is half-maximal. The exponent $n_{glc}$ reflects the cooperativity of glucose binding to the transporter, with higher values indicating greater cooperative interaction among glucose molecules. This equation captures the dependence of glucose transport rate on extracellular glucose concentration and the intrinsic properties of the GLUT1 transporter:

$$J_{GLUT} = g_{GLUT}\left(\frac{K_{glc}[Glc]_e}{M_{glc} + [Glc]_e}\right)^{n_{glc}}. \tag{11}$$

### 10. Glycolytic Model

To model the main chemical reaction described, we consider the conversion of glucose and ADP into pyruvate, ATP, NADH, and protons, represented by the equation:

$$\mathrm{Glucose} + 2\mathrm{NAD}^+ + 2\mathrm{ATP} + 2\mathrm{P_i} \underset{-}{\overset{+}{\rightleftharpoons}} 2\mathrm{Pyruvate} + 2\mathrm{NADH} + 2\mathrm{ATP} + 2\mathrm{H}^+.$$

The forward and backward reaction is given by:

$$F_{\mathrm{sGlyc}} = \left(\frac{[Glc]_c}{[Glc]_c + K_{glc}}\right)\left(\frac{[\mathrm{NAD}^+]_c}{[\mathrm{NAD}^+]_c + K_p}\right)^2\left(\frac{[\mathrm{ATP}]_c}{[\mathrm{ATP}]_c + K_{atp}}\right)^2\left(\frac{[\mathrm{P_i}]_c}{K_{P_i} + [\mathrm{P_i}]_c}\right)^2,$$

$$B_{\mathrm{Glyc}} = \left(\frac{[\mathrm{Pyr}]_c}{M_{cm_{pyr}} + [\mathrm{Pyr}]_c}\right)^2\left(\frac{[\mathrm{NADH}]_c}{[\mathrm{NADH}]_c + K_n}\right)^2\left(\frac{[\mathrm{ATP}]_c}{[\mathrm{ATP}]_c + K_{atp}}\right)^2\left(\frac{[\mathrm{H}^+]_c}{K_{\mathrm{H}^+} + [\mathrm{H}^+]_c}\right)^2.$$

Thus our glycolytic model is given by:

$$J_{Glyc} = g_{GLYC}\bigg(F_{\mathrm{Glyc}} - B_{\mathrm{Glyc}}\bigg). \tag{12}$$

The equation incorporates multiple terms that describe the concentration-dependent activities of substrates and products involved in glycolysis. These terms capture the saturation kinetics of glucose uptake and the inhibitory effects exerted by pyruvate, ATP, NADH, and protons. Each term is expressed as a substrate concentration divided by the sum of the substrate concentration and a constant, dependent on cooperative binding effects. All cofactor rate parameters were taken from Salem [14], Zhou et al. [13]. The remaining parameters were calculated as per our own data through $^{13}$C-MFA.

### 11. Lactate Dehydrogenase and the MCT-4 Lactate Channel

We use a model based on Falkenberg and Jakobsson [10] but matched to the dynamics by Zhou et al. [13], Salem [14]. Unlike the previous studies, we focus on also understanding pH concentration gradients, as our focus is essentially cellular respiration. The lactate dehydrogenase and the MCT-4 transporter play a role in pH regulation along with the Nhe, Ae2, and carbonic anhydrases. Its flux depends on the intracellular concentrations of pyruvate, NADH, lactate, $NAD^+$, and $H^+$. The equation models the rate of the LDH-catalysed reaction that converts pyruvate to lactate and vice versa, taking into account the enzyme's affinity for its substrates and products. The term $g_{LDH}$ is a scaling factor that adjusts the overall reaction rate, proportional to the density of active LDH enzymes:

$$F_{\mathrm{LDH}} = \left(\frac{[\mathrm{Pyr}]_c}{[\mathrm{Pyr}]_c + M_{cm_{pyr}}}\right)^2 \left(\frac{[\mathrm{NADH}]_c}{[\mathrm{NADH}]_c + K_n}\right)^2 \left(\frac{[\mathrm{H}^+]_c}{K_{\mathrm{H}^+} + [\mathrm{H}^+]_c}\right)^2,$$

$$B_{\mathrm{Lac}} = \left(\frac{[\mathrm{NAD}^+]_c}{[\mathrm{NAD}^+]_c + K_p}\right)^2 \left(\frac{[\mathrm{Lac}]_c}{[\mathrm{Lac}]_c + K_{Lac}}\right)^2.$$

$$J_{LDH} = g_{LDH}\Bigg(F_{\mathrm{LDH}} - B_{\mathrm{LDH}}\Bigg). \tag{13}$$

The MCT-4 flux model represents the activity of the cell membrane lactate $H^+$ symporter. Its flux depends on the lactate and $H^+$ concentrations in the cytoplasm and the extracellular space, the number of binding sites, and the half-saturation constants. The model evaluates the product of two terms and the exchanger's conductance, which is proportional to the density of active membrane proteins. The contribution from the lactate concentrations and binding site properties is represented by the Michaelis-Menten terms. Additionally, as a simplification, we include the difference of the transporter in reverse, which renders the exchanger bidirectional:

$$F_{\mathrm{MCT4}} = \left(\frac{[\mathrm{Lac}]_c}{[\mathrm{Lac}]_c + K_{Lac}}\right)\left(\frac{[\mathrm{H}^+]_c}{K_{\mathrm{H}^+} + [\mathrm{H}^+]_c}\right),$$

$$B_{\mathrm{MCT4}} = \left(\frac{[\mathrm{Lac}]_e}{[\mathrm{Lac}]_e + K_{Lac_e}}\right)\left(\frac{[\mathrm{H}^+]_e}{K_{\mathrm{H}^+} + [\mathrm{H}^+]_e}\right).$$

$$J_{\mathrm{MCT4}} = g_{\mathrm{MCT4}}\Bigg(F_{\mathrm{MCT4}} - B_{\mathrm{MCT4}}\Bigg) \tag{14}$$

#### 12. Aspartate-Glutamate, Malate-aKG transporters

The malate-aspartate shuttle relies on two mitochondrial transporters from the solute carrier 25a family. These are the Slc25a12/13 which exchanges, in an electroneutral manner, mitochondrial aspartate for cytosolyc glutamate with a $H^+$ to balance the aminoacid's negative charge:

$$\mathrm{Aspartate}_m + \mathrm{Glutamate}_c + \mathrm{H^+}_c \underset{-}{\overset{+}{\rightleftharpoons}} \mathrm{Aspartate}_c + \mathrm{Glutamate}_m + \mathrm{H^+}_c.$$

Similarly, the Slc25a10/11 exchanges, in an electroneutral manner, mitochondrial $\alpha$-ketoglutarate for cytosolyc malate:

$$\mathrm{Malate}_c + \alpha\mathrm{KG}_m \underset{-}{\overset{+}{\rightleftharpoons}} \mathrm{Malate}_m + \alpha\mathrm{KG}_c.$$

To model these we use a generic transporter equation, devised by Falkenberg and Jakobsson [10]:

$$J_{\mathrm{tr}} = G_{\mathrm{tr}}\left(\prod \frac{[\mathrm{met_i}]_c}{K_{\mathrm{met}} + [\mathrm{met_i}]_m} - \prod \frac{[\mathrm{met_j}]_c}{K_{\mathrm{met}} + [\mathrm{met_j}]_m}\right). \tag{15}$$

Here $k$ and $j$ represent the metabolite species exhanged across compartments. $G_{\mathrm{tr}}$ is the exchanger's membrane density and $K_{\mathrm{met}}$ the half-maximal activation of the transporter.

#### 13. Proton Motive Force

The Proton Motive Force (PMF) refers to the potential energy stored across a membrane due to the differential concentration of protons ($H^+$) on either side of that membrane. In the context of mitochondria, it's the difference in proton concentration across the inner mitochondrial membrane. This force is crucial for cellular energy production. Specifically, in the mitochondria, the electron transport chain creates a proton gradient, pumping protons from the matrix into the intermembrane space. This creates both a concentration gradient and an electrochemical gradient (or membrane potential difference).

The PMF is modelled as:

$$\Delta G_H = F\Delta\Psi + RT\ln\left(\frac{H_i}{H_m}\right). \tag{16}$$

The equation above is composed of two critical terms. Firstly, the electrochemical gradient is given by $F\Delta\Psi$, where $\Delta\Psi$ represents thee difference in membrane pootential across the inner mitochondrial membrane, $F$ is Faraday's constant in SI units. Similarly, the second term, $RT\ln\left(\frac{H_i}{H_m}\right)$ involves the proton concentration in the intermembrane space, and the concentration in the mitochondrial matrix. $R$ is the universal gas constant and $T$ the temperature. Adding both terms gives us Gibbs free energy change (or potential energy) due to the proton motove force. In other words, this equation quantifies the energy stored due to the difference in proton concentration and electrical potential across the compartments of the mitochondrion. This energy stored is used by the ATP synthase to synthesise ATP from ADP and phosphate.

### 14. ETC complex 1

Complex I, also known as NADH:ubiquinone oxidoreductase, is the first complex in the mitochondrial electron transport chain. It catalyses the transfer of electrons from NADH to ubiquinone ($Q$), resulting in the reduction of ubiquinone to ubiquinol ($QH_2$). During this process, protons are pumped from the matrix to the intermembrane space (IM space) of the mitochondria.

The model, created by Korzeniewski and Zoladz [15], sets the transporter's flux to be proportional to the difference in concentrations of NADH and $NAD^+$; given these drive the reaction, but its activity is modulated by an exponential term which is influenced by the Gibbs' free energy change associated with the process of pumping protons from one compartment to the other:

$$J_{\mathrm{C}_1} = {}_xC_1 \times \left( exp\left(\frac{-\Delta G_{C_1op} + 4\Delta G_H}{RT}\right) \times [\mathrm{NADH}]_m - [\mathrm{NAD}^+]_m \right), \quad (17)$$

where, the energy requuired, or released by the reaction is captured by the free energy change terms: $\Delta G_{C_1op}$ and $\Delta G_H$. $\Delta G_{C_1op}$ represents the energy difference when Complex 1 transfers electrons. The term incompasses three key principles: The standard free energy for the reaction, a term reepresenting the influence of proton concentration in the mitochondrion matrix along with the proton concentration at neutral mitochondrial (approx. $pH = 7$), and lastly, a term describing the ratio of ubiquinone to ubiquinol. As this last ratio changes, the energy change associated with the transfer of electrons varies. Thus:

$$\Delta G_{C_1op} = \Delta G_{C_10} - RT \ln \left( \frac{\mathrm{H}_m^+}{1e^{-7}} \right) - RT \ln \left( \frac{[\mathrm{Q}]_m}{[\mathrm{QH}_2]_m} \right),$$

$\Delta G_H$ is the proton motive force as described in C.1.

The model captures the essence of how the flux of protons $\mathrm{H}^+$ through Complex I varies. This flux is driven by the difference in NADH and $\mathrm{NAD}^+$ concentrations and is modulated by both the energy change associated with the electron transfer in Complex I and the proton pumping mechanism. It considers how energy efficient the process is, how much NADH is available, and the energy needed to pump protons across the mitochondrial membrane.

### 15. Electron Complex III

Complex III of the electron transport chain is responsible for transferring electrons from ubiquinol to cytochrome c. This model was constructed by Korzeniewski and Zoladz [15]. During the respiratory process, like C1, it acts as a proton pump across the inner mitochondrial membrane. Hence, similar to CI. Its flux thus, is quantifed by:

$$J_{C_3} = {}_xC_3\left(\frac{\frac{1+[\mathrm{Pi}]_m}{k_{Pi_3}}}{\frac{1+[\mathrm{Pi}]_m}{k_{Pi_4}}}\right)\times$$

$$\ldots\left(exp\left(\frac{-\left(\Delta G_{C_3op}+4\Delta G_H-2F\Delta\Psi_m\right)}{2RT}\right)[\mathrm{cytoC_{ox}}]_i-[\mathrm{cytoC_{red}}]_i\right). \tag{18}$$

CII's flux aims to capture the kinetics and thermodynamics of the proton pumping process by Complex III. Importantly, the equation is equuipped with a nuumerical factor indicating that inorganic phosphate plays a critical modulating role in the activity of this complex. This numerical factor, and the underlying biology it suggests proves a critical modelling angle in understanding mitochondrial swelling as we will see further. Moreover, the inclusion of phosphate as a modulating factor aligns with the reality of cellular metabolism. Phosphate plays critical roles in energy storage (ATP) and many enzymatic reactions. Modulation by phosphate can serve as a feedback mechanism to tune respiratory activity in response to cellular energy needs [16].

The free energy change in Complex III is mathematically represented by:

$$\Delta G_{C_3op} = \Delta G_{C_30} + 2RT\ln\left(\frac{[\mathrm{H}^+]_\mathrm{m}}{1e^{-7}}\right) - RT\ln\left(\frac{[\mathrm{QH_2}]_m}{[Q]_m}\right).$$

$\Delta G_{C_3op}$ can be dissected into three integral components: Firstly, it represents the standard free energy associated with reactions occurring within Complex III. Secondly, there is a component that accounts for the proton concentration within the mitochondrial matrix, denoted as

$$\frac{[\mathrm{H}^+]_\mathrm{m}}{1e^{-7}}.$$

Finally, a pivotal aspect of this equation is the term which signifies the ratio of the reduced state of ubiquinol to its counterpart, the oxidised form known as ubiquinone:

$$\frac{[\mathrm{QH_2}]_m}{[Q]_m}.$$

### 16. Electron Complex IV

Just as Compexes I and III, Complex IV's mechanistic proton pumping lies a process that culminates in the transfer of four distinct charges:

$$J_{C_4} = {}_xC_4\left(\frac{1+k_{O2}}{[\mathrm{O_2}]}\right)^{-1} \times \left(\frac{[\mathrm{cytoC_{red}}]_m}{[\mathrm{cytoC_{tot}}]_i}\right) \times ...$$
$$\left(exp\left(\frac{-\left(\Delta G_{C_4op} + 2\Delta G_H\right)}{2RT}\right) \times ...\right.$$
$$\left.[\mathrm{cytoC_{red}}]_m - [\mathrm{cytoC_{ox}}]_m \times exp\left(\frac{F\Delta\Psi}{RT}\right)\right). \tag{19}$$

The Complex IV flux is expressed in a form similar to those of complexes I and III, with a few key differences. The model is equipped with a dependency on oxygen concentration. Its presence suggests that the reaction kinetics may not be linear with respect to the oxygen concentration and may be modified by this factor. The term is also equipped with a proportion of reduced cytochrome c in the matrix relative to the total amount available. As the amount of reduced cytochrome c increases, the flux through Complex IV will also likely increase since reduced cytochrome c is a substrate for the reaction. Just as complexes I and III, the exponential components are essentially accounting for the thermodynamic driving forces of the reaction:

$$\Delta G_{C_4op} = \Delta G_{C_40} - 2RT\ln\left(\frac{[\mathrm{H^+}]_m}{1e^{-7}}\right) - \frac{RT}{2}\ln\left([\mathrm{O_2}]\right).$$

Here, the term with captures the free energy change associated with the process in Complex IV and the proton motive energy, respectively. Dividing RT by 2 denotes how it quantifies movement per-electron basis, given that the overall reaction transfers two electrons. Finally, cytochrome c Redox state. This difference reflects the net availability of reduced cytochrome c for the reaction. The magnitude and direction of this difference can drive the flux either forward or backward, depending on whether there's more reduced or oxidised cytochrome c.

### 17. ATP Synthase

ATP Synthase is responsible for converting ADP (Adenosine Diphosphate) into ATP (Adenosine Triphosphate) within the mitochondrial matrix. This conversion is a pivotal step in cellular energy production. The process by which this transformation occurs is modelled using the flux equation for ATP synthase, denoted as:

$$J_{F_1} = {}_xF_1 \times \left( exp\left( \frac{-(\Delta G_{F_10} - n_A \Delta G_H)}{RT} \right) \times \dots \right.$$
$$\left. \left( \frac{K_{DD}}{K_{DT}} \right) \times \left[^{\mathrm{Mg}^{2+}}\mathrm{ADP}\right]_m [\mathrm{Pi}]_m - \left[^{\mathrm{Mg}^{2+}}\mathrm{ATP}\right]_m \right). \quad (20)$$

The flux equation involves, much as CI - CIV, the standard free energy for the conversion of ADP to ATP. The quantity denotes the free energy contribution from proton transport, this time from intermembrane space to the mitochondrion matrix. Specifically, it accounts for the number of protons that are moved across the mitochondrial inner membrane for each conversion of ADP to ATP (which is approximately $n_A = 3$) , thereby influencing the reaction's energetics. The negative sign in the exponential ensures that the reaction is energetically favourable. The model also features a ratio of equilibrium constants or binding affinities, representing the ratio between ADP dissociation to ATP dissociation aspect of the reaction mechanism. Fuurthermore, explicitly included, are the matrix concentrations of magnesium-bound ($Mg^{2+}$) ADP, inorganic phosphate, and magnesium-bound ATP, respectively [17]. The reaction's forward rate depends on the availability of magnesium-bound ADP and inorganic phosphate, while its reverse rate is influenced by the concentration of magnesium-bound ATP. Their incluusion capitulates a more accurate description of the process, and also serve as cofactors in the phosphorylation process. The entire expression is takes into account the energy requirements and yields, ensuring that energy is conserved and that the flux aligns with the principles of thermodynamics.

### 18. Adenine nucleotide translocator (ANT)

ANT flux involves the displacement of one negative charge from the matrix to the motochondrial inner-membrane space, and is therefore coupled to the electrostatic membrane potential [18, 15, 19, 20]; the following empirical expression is used to model the ANT flux:

$$J_{\mathrm{ANT}} = X_{\mathrm{ANT}} \left( \frac{[\mathrm{fADP}]_i}{[\mathrm{fADP}]_i + [\mathrm{fATP}]_i e^{-0.35F\Delta\Psi/RT}} \right. \\ \left( \frac{[\mathrm{fADP}]_m}{[\mathrm{fADP}]_m + [\mathrm{fATP}]_m e^{0.65F\Delta\Psi/RT}} \right) \\ \left. \left( \frac{1}{1 + k_{m,\mathrm{ADP}}/[\mathrm{fADP}]_i} \right) \right). \tag{21}$$

The flux model assumes the adeninde nucleotide translocator operates on $Mg^{2+}$-unbound ATP and ADP in the two compartments [15].

### 19. Magnesium Ion Binding to ATP

Magnesium ($Mg^{2+}$) is pivotal in the biochemistry of ATP. Comprising a ribose sugar, adenine, and three phosphate groups, ATP's phosphate groups carry a negative charge, confer on the molecule with a pronounced negative charge density [17, 21]. $Mg^{2+}$ form bonds with these phosphate groups, effectively neutralising some of the inherent repulsion between them, thereby stabilising ATP. A host of enzymes, such as kinases, that engage with ATP have binding sites specifically crafted to fit the ATP-$Mg^{2+}$ complex. This dictates that for optimal enzyme functionality, ATP should be complexed with magnesium [17, 21]. The $Mg^{2+}$ presence aids in the precise positioning of ATP within these enzymes' active sites, enhancing the efficiency of catalysis. When ATP undergoes hydrolysis to ADP and an inorganic phosphate, it releases energy in an exergonic reaction. Magnesium is instrumental in the appropriate coordination of the phosphates during this process. In a water-based environment, ATP might undergo spontaneous hydrolysis, resulting in an energy storage loss. By binding with ATP, magnesium reduces the rate of this involuntary hydrolysis, further stabilising the ATP molecule. This magnesium binding triggers specific conformational alterations in ATP, crucial for its synergy with various proteins and enzymes. In essence, magnesium's role in ATP's biochemistry is multifaceted, influencing its stability, interactions, and its esteemed position as the cellular energy currency [22, 16, 17].

Binding between magnesium ion and ATP and ADP is driven via the following fluxes:

$$J_{\text{MgATP}_m} = X_{MgA}\left([\text{fATP}]_m[\text{Mg}^{2+}]_m - K_{\text{MgATP}}[\text{mATP}]_m\right), \tag{22}$$

$$J_{\text{MgADP}_m} = X_{MgA}\left([\text{fADP}]_m[\text{Mg}^{2+}]_m - K_{\text{MgADP}}[\text{mADP}]_m\right), \tag{23}$$

$$J_{\text{MgATP}_i} = X_{MgA}\left([\text{fATP}]_i[\text{Mg}^{2+}]_i - K_{\text{MgATP}}[\text{mATP}]_i\right), \tag{24}$$

$$J_{\text{MgADP}_i} = X_{MgA}\left([\text{fADP}]_i[\text{Mg}^{2+}]_i - K_{\text{MgADP}}[\text{mADP}]_i\right). \tag{25}$$

Notice how in the equations above, the unbond ATP and ADP concentrations are denoted: [fATP] and [fADP]. They are free ($f$), as they are not bound to $Mg^{2+}$ in the mitochondrial matrix and its inter-membrane space. Similarly, we denote [mATP] and [mADP] the magnesium-bound ATP and ADP concentrations in the mitochondrial matrix and its inter-membrane space. The parameter $X_{MgA}$ is the forward binding rate constant for the binding reactions; the effective unbinding constant is computed to satisfy the equilibrium dissociation relations for ATP and ADP binding with $Mg^{2+}$.

### 20. The mitochondrial phosphate carrier (PiC - Slc25a3)

The mitochondrial phosphate carrier (PiC) stands as a solute carrier protein within the mitochondria, and its genetic blueprint is encoded by the solute carrier family 25a3 (Slc25a3) in humans [23, 24]. Acting as a pivotal courier, PiC ferries phosphate — an essential substrate for oxidative phosphorylation — through the inner mitochondrial membrane. Beyond this primary role, its transport activities play a consequential part in facilitating efficient mitochondrial calcium management [23, 24]. Transport of inorganic phosphate between the matrix and IM space is coupled to the hydrogen ion gradient. It is assumed that $H^+$ and $H_2PO_4$ are transported by a cotransport process, with $H^+$ and $H_2PO_4$ moving together across the membrane in a 1:1 ratio in a net electroneutral exchange. We assume $H^+$ binding to Pi is assumed to be in equilibrium on either side of the membrane with,

$$[\mathrm{H_2PO_4^-}]_i = \frac{[\mathrm{H}^+]_i[\mathrm{Pi}]_i}{\Big([\mathrm{H}^+]_i + k_{dH}\Big)}, \text{ and}$$
$$[\mathrm{H_2PO_4^-}]_m = \frac{[\mathrm{H}^+]_m[\mathrm{Pi}]_m}{\Big([\mathrm{H}^+]_m + k_{dH}\Big)},$$

where, $k_{dH}$ is the dissociation constant for the reaction. In these expressions Pi represents the sum of species $\mathrm{H_2PO_4^-}$ and $\mathrm{H_2PO_4^{2-}}$. The PiC flux, then, is modeled as reversible Michaelis–Menten flux:

$$J_{\mathrm{PiC}} = X_{\mathrm{PiC}} \frac{\Bigg(\mathrm{H}_m^+[\mathrm{H_2PO_4^-}]_i - \mathrm{H}_i^+[\mathrm{H_2PO_4^-}]_m\Bigg)}{\Bigg([\mathrm{H_2PO_4^-}]_i + k_{\mathrm{PiH}}\Bigg)}. \tag{26}$$

Here $X_{\mathrm{PiC}}$ steady-state imposed parameter and $k_{\mathrm{PiH}}$ is the Michaelis–Menten constant for $\mathrm{H_2PO_4^-}$ on the outside of the membrane.

### 21. Adenyl Kinase

Adenylate kinase (AK) is a ubiquitous enzyme that contributes to the homeostasis of the cellular adenine nucleotide composition. Three isozymes, AK1, AK2, and AK3, have so far been characterized in vertebrates. They are located in different tissues, while their primary and tertiary structures are similar [25, 26]. In this model, the flux of high-energy phosphates are co-transferred between ATP, ADP, and AMP in the inner-membrane space via a general AK reaction; regardless of the isozymes [25, 26]:

$$J_{\mathrm{AK}} = X_{\mathrm{AK}} \Bigg( K_{\mathrm{AK}}[\mathrm{ADP}]_i[\mathrm{ADP}]_i - [\mathrm{AMP}]_i[\mathrm{ATP}]_i \Bigg), \tag{27}$$

where $K_{\mathrm{AK}} = 0.4331$ is the equilibrium constant for the reaction, and $X_{\mathrm{AK}}$ is the AK enzyme activity parameter.

### 22. Proton leak and potassium fluxes

The present work assumes that $Ca^{2+}$ concentrations and fluxes have only secondary effects on membrane potential compared to the primary effects of currents associated with the respiratory chain, the ANT current, and the proton leak. Therefore, fluxes of calcium are not considered at this stage. However, they will be integrated in further iterations oof the model. Since $K^+$ is required to buffer the matrix pH and $Mg^{2+}$ is required for ATP synthesis and the ANT flux, these ions are considered in the model. These are modelled using the Goldman-Katz-Hutchkin equation (obtained by solving the one-dimensional Nernst–Planck equation) [9]:

$$J_{Hle} = X_{\mathrm{Hlc}} \Delta\Psi \frac{\left( \mathrm{H}_i^+ \; e^{(F\Delta\Psi/RT)} - [H^+]_m \right)}{\left( e^{(F\Delta\Psi/RT)} - 1 \right)}, \tag{28}$$

and

$$J_K = X_K \Delta\Psi \frac{\left( [\mathrm{K}^+]_i e^{(F\Delta\Psi/RT)} - [\mathrm{K}^+]_m \right)}{\left( e^{(F\Delta\Psi/RT)} - 1 \right)}. \tag{29}$$

### 23. Potassium/Proton exchanger

The mitochondrial potassium/proton exchanger is an integral component of the mitochondrial inner membrane that facilitates the movement of potassium ions into the mitochondrial matrix in exchange for protons. This exchanger plays a crucial role in maintaining the mitochondrial membrane potential and pH gradient, which are essential for the production of ATP through oxidative phosphorylation. The proper functioning of this exchanger is pivotal for cellular energy metabolism and overall cell viability. Disruptions or abnormalities in its activity can have profound implications on cellular health and have been implicated in various pathological conditions [27]. In this model we use a linear exchange model as described by (Keener and Sneyd 2010)[9]:

$$J_{KH} = X_{KH}\left([\mathrm{K}^+]_i \mathrm{H}_m^+ - [\mathrm{K}^+]_m \mathrm{H}_i^+\right). \tag{30}$$

### 24. AMP ATP and ADP

Transport of ATP, ADP, AMP, and Pi between the cytosol and the mitochondrial inter-membrane space is modelled using linear transfer between compartments. The models can be changed to account for a more physiological representation; however, in this model the scope is to have a detailed enough mechanism to explore SDH, and not metabolite transport. Thus, for simplicity sake, the transport is modelled according to the following fluxes:

$$J_{\mathrm{ATP}} = \gamma p_A\left([\mathrm{ATP}]_e - [\mathrm{ATP}]_i\right), \tag{31}$$

$$J_{\mathrm{ADP}} = \gamma p_A\left([\mathrm{ADP}]_e - [\mathrm{ADP}]_i\right), \tag{32}$$

$$J_{\mathrm{AMP}} = \gamma p_A\left([\mathrm{AMP}]_e - [\mathrm{AMP}]_i\right), \tag{33}$$

$$J_{\mathrm{Pi}} = \gamma p_{Pi}([\mathrm{Pi}]_e - [\mathrm{Pi}]_i). \tag{34}$$

These equations denote the transport of ATP, ADP, AMP, and Pi from the cytosol to the inter-membrane space. Each flux term is modelled as a linear flux proportional to the membrane permeability to each metabolite and the compartments volume ratio (i.e., mitochondrial inner-membrane to cytosol).

### 25. Succinate Dehydrogenase - ETC Complex 2

### 26. Binding Polynomials for enzyme, substrates, products and regulators

A binding polynomial model, arises as the denominator of the rational function describing the average number of occupied binding sites ($K_d$ - parameter) as a function of ligand (substrate) activity [9, 28]. In other words, the binding polynomials represent the likelihood of different molecules binding to specific sites on the enzyme. The mathematical representation of the fraction of enzyme with a specific ligand bound to it, is as follows:

$$BP_Q = 1 + \frac{[\mathrm{Q}]_\mathrm{i}}{K_{d_Q}} + \frac{[\mathrm{QH_2}]_i}{K_{d_{QH_2}}}, \tag{35}$$

$$BP_{\mathrm{FAD^+}} = 1 + \frac{[\mathrm{succ}]_m}{K_{d_\mathrm{SUCC}}} + \frac{[\mathrm{fum}]_m}{K_{d_\mathrm{FUM}}}. \tag{36}$$

In Eqs. 35 and 36, it is possible to observe that the relationship between the ligand's activity (how active or effective it is in binding) and the average number of occupied sites can be described as a rational function. A rational function is a ratio of two polynomials. In this context, the denominator of this function is particularly noteworthy as it represents the binding polynomial [9]. Moreover, the dissociation constant, $K_d$ is a specific parameter that gives an idea about the affinity between a ligand and its binding site. Thus, Eqs. 35 and 36 offer a mathematical way to quantify the behaviour of ligands as they bind to their target molecules. The model looks at how the average number of occupied binding sites changes with varying ligand activity, encapsulating this relationship within the framework of a rational function. The denominator of this function, representing the binding polynomial, is closely tied to the ligand's dissociation constant ($K_d$) providing insights into the binding affinity [9].

### 27. Midpoint potential pH corrections

Redox centres are regions of proteins (often metal ions or organic compounds) that can transfer electrons between molecules. In the case of the succinate deehydrogenase these are the iron-sulfur clusters in the SDH$_b$ subunit. The formation energy describes the energy change when these centers are formed.

In the model we calculate the midpoint potential,$E_m$, (i.e., the potential at which half of the molecules of a particular redox couple are in the oxidised state and half are in the reduced state) of the SDH redox centres. In other words, at this potential, the concentrations of the oxidised and reduced species are equal. It serves as an intrinsic property of a redox couple, effectively indicating its tendency to either lose or gain electrons [9]. Although this model serves as a reference point to understand the tendency of a molecule to accept or donate electrons under stadard conditions, the purpose of our model is to underrstand actual conditions. Thus, we use the Nernst potentials in combination with midpoint potentials. In this way our model can account for influences such as the pH of the surrounding environment. This is especially relevant for redox reactions involving protons. As pH changes, it can shift the balance between the oxidised and reduced states, effectively altering the midpoint potential [9]. This model serves as the theoretical foundation for the necessary adjustment, linking changes in potential to shifts in pH and other conditions. Flavins', i.e., $\mathrm{FAD}^+ \rightarrow \mathrm{FADH} \rightarrow \mathrm{FADH}_2$, willingness to donate electrons, (midpoint potentials), are modelled as:

$$E_{m_{\mathrm{FAD}^+\rightarrow\mathrm{FADH}}} = E_{m0_{\mathrm{FAD}^+\rightarrow\mathrm{FADH}}} + \frac{RT}{F} \ln \left( \mathrm{H}_m^+ \; \mathrm{H}_{i_{\mathrm{FADH}}} \right), \tag{37}$$

$$E_{m_{\mathrm{FADH}\rightarrow\mathrm{FADH}_2}} = E_{m0_{\mathrm{FADH}\rightarrow\mathrm{FADH}_2}} + \frac{RT}{F} \ln \left( \mathrm{H}_m^+ \frac{\mathrm{H}_{\mathrm{FADH}_2}}{\mathrm{H}_{\mathrm{FADH}}} \right), \tag{38}$$

$$E_{m_{\mathrm{FAD}^+\rightarrow\mathrm{FADH}_2}} = E_{m0_{\mathrm{FAD}^+\rightarrow\mathrm{FADH}_2}} + \frac{RT}{F} \ln \left( (\mathrm{H}_m^+)^2 \frac{\mathrm{H}_{\mathrm{FADH}_2}\mathrm{H}_{\mathrm{iFADH}}}{\mathrm{H}_{\mathrm{FADH}}} \right), \tag{39}$$

where,

$$\mathrm{H}_{\mathrm{FADH}} = 1 + \frac{\mathrm{H}_m^+}{\mathrm{K}_{\mathrm{FADH}}},$$
$$\mathrm{H}_{\mathrm{FADH}_2} = 1 + \frac{\mathrm{H}_m^+}{K_{\mathrm{FADH}_2}},$$
$$\mathrm{H}_{\mathrm{iFADH}} = 1 + \frac{\mathrm{K}_{\mathrm{FADH}}}{\mathrm{H}_m^+}.$$

The reduction of ubiquinon to ubiquinol ($Q$ to $QH_2$, respectively), is also equipped midpoint potential term, where an intermediary state ($SQ$) is added for modelling purposes (where only one electron is donated) as follows:

$$\begin{aligned}
E_{m_{Q\to SQ}} &= E_{m0_{Q\to SQ}}, \\
E_{m_{SQ\to QH_2}} &= 2E_{m0_{SQ\to QH_2}} + 2\frac{RT}{F}\ln\left(\mathrm{H}_m^+\right) - E_{m_{Q\to SQ}}, \\
E_{m_{Q\to QH_2}} &= E_{m0_{Q\to QH_2}} + \frac{RT}{F}\ln\left(\mathrm{H}_m^+\right).
\end{aligned}$$

The corrected redox midpoint potentials for the TCA's succinate to fumarate are given by:

$$E_{m_{\mathrm{FUM}\to\mathrm{SUCC}}} = E_{m0_{\mathrm{FUM}\to\mathrm{SUCC}}} + \frac{RT}{F}\ln\left(\mathrm{H}_m^+\right).$$

Finally, we define the midpoint potentials for electron transfer of bound states as:

$$E_{mb_{Q\to SQ}} = E_{m_{Q\to SQ}} + \frac{RT}{F}\ln\left(K_{d_Q}\right), \tag{40}$$

$$E_{mb_{SQ\to QH_2}} = E_{m_{SQ\to QH_2}} - \frac{RT}{F}\ln\left(Kd_{QH_2}\right), \tag{41}$$

$$E_{mb_{\mathrm{SUCC}\to\mathrm{FUM}}} = E_{m_{\mathrm{FUM}\to\mathrm{SUCC}}} - \frac{RT}{F2}\ln\left(K_{d_{\mathrm{SUC}}}\right) + \frac{RT}{F2}\ln\left(K_{d_{\mathrm{FUM}}}\right). \tag{42}$$

Eqs. 37 to 42 account for how metabolite species give away or accept electrons (and thus undergo a redox reaction) might change as the pH of the mitochondria changes.

*27.1. Boltzman redox poise potentials*

The following modelling approach aims to understand the energetics of the redox processes occuring in SDH. Firstly, by converting midpoint potentials into Gibbs free energies, we're able to gain insights into the thermodynamic feasibility of reactions and the energetic interactions between substrates and redox centers:

$$\Delta G = -nF * \Delta E_h,$$

where $n$ is the number of electrons transferred in the reaction, and $E_h$ are the Boltzman redox poise potentials (given in Appendix). Thus, to determine the transition rates, each combination of the redox centres reduced in each electrinic state are calculated by Boltzmann distributions. We define $S_r^k$ the fraction of redox centres $r$ existing in the electrinic state $k$ that is reduced, and $\Delta G_r^k$, are the free energy changes for each redox centre $r$ calculateed from the linear superposition of the midpoint potentials:

$$S_r^k = \frac{e^{\left(-\Delta G_r^k/RT\right)}}{\sum_r e^{\left(-\Delta G_r^k/RT\right)}}. \tag{43}$$

In Eq. 43, $S_r^k$ is the fraction of redox centres $r$ existing in the oxidation state $k$ that is reduced, and $\Delta G_r^k$ is the free energy change for each redox centre $r$ calculated from the linear super position of adjusted midpoint potentials (Eqs. 37 to 42). To calculate the free energy change for the combination of redox centers, the individual free energies for the redox center reactions are summed (i.e., they are independent from each other).

#### *27.2. State Transitions*

As the mathematical sub-model of the SDH is geared towards modelling the redox state of the enzyme, we utilise a modified version of a five redox state transition kinetic model developed by (Neeraj et al., 2020) [29]. This model describes the potential redox states ($E_i$) of the enzyme, where '$i$' represents the redox state correlating to the total number of electrons present on the enzyme. The transition rates between these redox states are denoted as '$k_i^j$'. Pertinent redox reactions underpinning the redox state transitions are mediated by either one or two electrons. The edges connecting the oxidation states depicted in [FIGURE] represent the partial reactions governing the connection between state $i$ and state $j$. These reaction rates, '$k_i^j$', embody molecular processes such as the reduction of FAD to $FADH_2$ by succinate, the oxidation of FADH or $FADH_2$ by oxygen, and Q reduction at the Q site. Before the enzyme can transition between oxidation states, it must adopt the appropriate enzyme-substrate complex configuration. For instance, the oxidation of succinate can only take place when the FAD-binding site is ready for succinate binding and the FAD is in a fully oxidised state. Binding polynomials are employed to determine the proportion of succinate bound to the complex, whilst the Boltzmann distribution calculates the fraction of the protein complex with a fully oxidised FAD within a given oxidation state. The net steady-state rate of succinate oxidation is then determined by totalling the oxidation state transition rates. The state transition equations are fully described in the supporting material of this article.

The resulting fluxes for the overal SDH reaction - Succinate + FAD + Q $\leftrightarrow$ Fumarate + $QH_2$ + $FADH_2$, are given by:

$$
\begin{aligned}
J_{\mathrm{succ}} = E_{tot}\Bigg( & E_0 k_{02_{\mathrm{SUCC}\to\mathrm{FAD}}} + E_1 k_{13_{\mathrm{SUCC}\to\mathrm{FAD}}} \\
& + E_2 k_{24_{\mathrm{SUCC}\to\mathrm{FAD}}} - E_2 k_{20_{\mathrm{FUM}\to\mathrm{FADH_2}}} \\
& - E_3 k_{31_{\mathrm{FUM}\to\mathrm{FADH_2}}} - E_4 k_{42_{\mathrm{FUM}\to\mathrm{FADH_2}}} \Bigg),
\end{aligned}
\tag{44}
$$

$$
\begin{aligned}
J_{\mathrm{QH_2}} = E_{tot}\Bigg( & E_2 k_{20_{\mathrm{SQ-ISC_3}\to\mathrm{QH_2}}} + E_3 k_{31_{\mathrm{SQ-ISC_3}\to\mathrm{QH_2}}} \\
& + E_4 k_{42_{\mathrm{SQ-ISC_3}\to\mathrm{QH_2}}} - E_0 k_{02_{\mathrm{QH_2}\to\mathrm{SQ-ISC_{3_{ox}}}}} \\
& - E_1 k_{13_{\mathrm{QH_2}\to\mathrm{SQ-ISC_{3_{ox}}}}} - E_2 k_{24_{\mathrm{QH_2}\to\mathrm{SQ-ISC_{3_{ox}}}}} \Bigg).
\end{aligned}
\tag{45}
$$

### 28. Model Equations

Our ordinary differential equation model takes on the following form:

$$\omega_c \frac{\mathrm{d}[\mathrm{Glc}]_c}{\mathrm{d}t} = J_{Glut} - J_{Glyc}, \tag{46}$$

$$\omega_c \frac{\mathrm{d}[\mathrm{Pyr}]_c}{\mathrm{d}t} = 2\Bigg( J_{Glyc} - J_{Ldh} - J_{MPC} \Bigg), \tag{47}$$

$$\omega_c \frac{\mathrm{d}[\mathrm{Lac}]_c}{\mathrm{d}t} = 2\Bigg( J_{Ldh} - J_{MCT4} \Bigg), \tag{48}$$

$$\omega_c \frac{\mathrm{d}[\mathrm{NADH}]_c}{\mathrm{d}t} = 2\Bigg( J_{Glyc} - J_{Ldh} \Bigg) - J_{MDH_c}, \tag{49}$$

$$\omega_c \frac{\mathrm{d}[\mathrm{ATP}]_c}{\mathrm{d}t} = 2J_{Glyc} - J_{ATP} + J_{ANT}, \tag{50}$$

$$\omega_c \frac{\mathrm{d}[\mathrm{ADP}]_c}{\mathrm{d}t} = J_{ATP} - 2J_{Glyc} - J_{ANT}, \tag{51}$$

$$\omega_c \frac{\mathrm{d}[\mathrm{H}^+]_c}{\mathrm{d}t} = 2\Bigg( J_{Glyc} - J_{MPC} - J_{Ldh} \Bigg) + J_{atp} - J_{Nhe} + J_{CA}, \tag{52}$$

$$\omega_c \frac{\mathrm{d}[\mathrm{Asp}]_c}{\mathrm{d}t} = J_{aspglu} - J_{Aaspta_c}, \tag{53}$$

$$\omega_c \frac{\mathrm{d}[\mathrm{Oaa}]_c}{\mathrm{d}t} = J_{Aaspta_c} - J_{MDH_c}, \tag{54}$$

$$\omega_c \frac{\mathrm{d}[\mathrm{Mal}]_c}{\mathrm{d}t} = J_{MDH_c} - J_{\alpha kgmal}, \tag{55}$$

$$\omega_c \frac{\mathrm{d}[\mathrm{Glu}]_c}{\mathrm{d}t} = J_{Glu} - J_{Aaspta_c} - J_{Glu_m}, \tag{56}$$

$$\omega_c \frac{\mathrm{d}[\mathrm{Cl}^-]_c}{\mathrm{d}t} = J_{Ae2} - 2J_{Nkcc} - J_{Cl}, \tag{57}$$

$$\omega_c \frac{\mathrm{d}[\mathrm{K}^+]_c}{\mathrm{d}t} = J_{Nkcc} + 2J_{NaK} - J_{K}, \tag{58}$$

$$\omega_c \frac{\mathrm{d}[\mathrm{Na}^+]_c}{\mathrm{d}t} = J_{Nhe} - 3J_{NaK} + J_{Nkcc}, \tag{59}$$

$$\omega_c \frac{\mathrm{d}[\mathrm{HCO}_3^+]_c}{\mathrm{d}t} = J_{CA} - J_{Ae2}, \tag{60}$$

$$C_{m_c} \frac{\mathrm{d}V_m}{\mathrm{d}t} = J_{Cl} + J_{K} - J_{NaK}. \tag{61}$$

$$\omega_m \frac{\mathrm{d}[\mathrm{Pyr}]_m}{\mathrm{d}t} = 2J_{MPC} - J_{PDH} - J_{PC}, \tag{62}$$

$$\omega_m \frac{\mathrm{d}[\mathrm{NADH}]_m}{\mathrm{d}t} = J_{mdh_c} + J_{\alpha kgd} + J_{PDH} - J_{C_1}, \tag{63}$$

$$\omega_m \frac{\mathrm{d}[\mathrm{aCoA}]_m}{\mathrm{d}t} = J_{PDH} - J_{CSm}, \tag{64}$$

$$\omega_m \frac{\mathrm{d}[\mathrm{H}^+]_m}{\mathrm{d}t} = 2\left(J_{MPC} - J_{C_3}\right) + J_{CSm} + 3J_{ATPS} - 4\left(J_{C_1} + J_{C_4}\right) + J_{Hle} + J_{KH}, \tag{65}$$

$$\omega_m \frac{\mathrm{d}[\mathrm{Cit}]_m}{\mathrm{d}t} = J_{CSm} - J_{icdh}, \tag{66}$$

$$\omega_m \frac{\mathrm{d}[\alpha\mathrm{KG}]_m}{\mathrm{d}t} = J_{icdh} - J_{\alpha kgd} - J_{\alpha kgmal}, \tag{67}$$

$$\omega_m \frac{\mathrm{d}[\mathrm{ATP}]_m}{\mathrm{d}t} = J_{\alpha kgd} + J_{ATPS} - J_{ANT}, \tag{68}$$

$$\omega_m \frac{\mathrm{d}[\mathrm{ADP}]_m}{\mathrm{d}t} = -\omega_m \frac{\mathrm{d}[\mathrm{ATP}]_m}{\mathrm{d}t}, \tag{69}$$

$$\omega_m \frac{\mathrm{d}[\mathrm{Succ}]_m}{\mathrm{d}t} = J_{\alpha kgd} - J_{suc} - J_{st}, \tag{70}$$

$$\omega_m \frac{\mathrm{d}[\mathrm{Fum}]_m}{\mathrm{d}t} = J_{suc} - J_{fum}, \tag{71}$$

$$\omega_m \frac{\mathrm{d}[\mathrm{Mal}]_m}{\mathrm{d}t} = J_{Fum} - J_{mdh_m} + J_{\alpha kgmal}, \tag{72}$$

$$\omega_m \frac{\mathrm{d}[\mathrm{Oaa}]_m}{\mathrm{d}t} = J_{mdh_m} - J_{CSm} - J_{Aaspta_m} + J_{PC}, \tag{73}$$

$$\omega_m \frac{\mathrm{d}[\mathrm{QH}_2]_m}{\mathrm{d}t} = J_{C_1} + J_{qh_2} - J_{C_3}, \tag{74}$$

$$\omega_m \frac{\mathrm{d}[\mathrm{Asp}]_m}{\mathrm{d}t} = J_{Aaspta_m} - J_{aspglu}, \tag{75}$$

$$\omega_m \frac{\mathrm{d}[\mathrm{Glu}]_m}{\mathrm{d}t} = J_{Glu_m} - J_{aspglu} \tag{76}$$

$$\omega_i \frac{\mathrm{d}[\mathrm{H}^+]_i}{\mathrm{d}t} = 2J_{C_3} - 3J_{ATPS} + 4\left(J_{C_1} + J_{C_4}\right) - J_{Hle} - J_{KH} - J_{P_iH}, \quad (77)$$

$$\omega_i \frac{\mathrm{d}[\mathrm{ATP}]_i}{\mathrm{d}t} = J_{ANT} + J_{ATPMg} - J_{ATP_t}, \quad (78)$$

$$\omega_i \frac{\mathrm{d}[\mathrm{ADP}]_i}{\mathrm{d}t} = -J_{ANT} + 2J_{ATPMg} + J_{ADP_t}, \quad (79)$$

$$\omega_i \frac{\mathrm{d}[\mathrm{Cred}]_i}{\mathrm{d}t} = 2J_{C_3} - 4J_{C_4}, \quad (80)$$

$$C_{m_i} \frac{\mathrm{d}\Delta\Psi}{\mathrm{d}t} = 4\left(J_{C_1} + J_{C_4}\right) + 2J_{C_3} - 3J_{ATPS} - J_{ANT} - J_{Hle} - J_K. \quad (81)$$

$$\frac{\mathrm{d}\omega_c}{\mathrm{d}t} = \sigma_c\left(\sum[\mathrm{i/m}]_c - [\mathrm{i/m}]_e - [\mathrm{i/m}]_m\right) \quad (82)$$

$$\frac{\mathrm{d}\omega_m}{\mathrm{d}t} = \sigma_m\left(\sum[\mathrm{i/m}]_m - [\mathrm{i/m}]_i - [\mathrm{i/m}]_c\right). \quad (83)$$

### 29. Levenberg-Marquardt Algorithm

The Levenberg-Marquardt algorithm is a numerical optimisation technique that combines the principles of both the gradient descent and the Gauss-Newton method [30, 31]. It is specifically designed to address non-linear least squares problems, making it particularly suitable for complex model fitting in scenarios where an optimal solution must be found in the context of curve-fitting and parameter estimation[30, 31].

The algorithm seeks to minimise a sum of squared function values (residuals), which quantify the discrepancy between observed data and model predictions based on a set of parameters. Update Rule: At each iteration, parameters are updated by solving the equation:

$$\mathbf{J}^T\mathbf{J} + \lambda\mathbf{D}\Delta\mathbf{p} = \mathbf{J}^T\mathbf{r}, \quad (84)$$

where $\mathbf{J}$ is the Jacobian matrix of partial derivatives of the residuals with respect to the parameters, $\mathbf{r}$ is the vector of residuals, $\lambda$ is the damping factor that adjusts the behavior of the algorithm, $\mathbf{D}$ is a diagonal matrix, typically taken as the diagonal of $\mathbf{J}^T\mathbf{J}$, $\Delta\mathbf{p}$ is the step to be taken in parameter space.

The damping factor $\lambda$ plays a crucial role in the convergence behavior of the algorithm. A larger $\lambda$ makes the algorithm behave more like a gradient descent (robust but slow), whereas a smaller $\lambda$ leads to a behavior closer to the Gauss-Newton method (fast but potentially unstable). The adjustment of $\lambda$ is based on the reduction of the residual: if the adjustment leads to a decrease in the residual, $\lambda$ is decreased, promoting faster convergence; if not, $\lambda$ is increased to gain stability.

The algorithm iterates until a stopping criterion is met, which could be based on the size of the gradient (norm of $\mathbf{J}^T\mathbf{r}$, the change in parameter values, or the improvement in the residual norm.

The Levenberg-Marquardt algorithm provides a robust and efficient method for dealing with a wide range of problems, and balances between the rapid convergence of the Gauss-Newton method and the stability of the gradient descent [30, 31].

### 30. Transporters

$$
\begin{aligned}
\frac{dx_1}{dt} &= k_1[P]_i x_2 + k_4 y_1 - x_1\Bigg(k_1[S]_i + k_4\Bigg),\\
\frac{dx_2}{dt} &= k_{-2} y_2 + k_1[S]_i x_1 - x_2\Bigg(k_2 + k_{-1}[P]_i\Bigg),\\
\frac{dy_1}{dt} &= k_{-4} x_1 + k_3[P]_e y_2 - y_1\Bigg(k_4 + k_{-3}[S]_e\Bigg),\\
x_1 &+ x_2 + y_1 + y_2 = 1.
\end{aligned}
$$

### 31. Steady-State Conditions

At steady state, the time derivatives are set to zero:

$$
\frac{dx_1}{dt} = 0,\quad
\frac{dx_2}{dt} = 0,\quad
\frac{dy_1}{dt} = 0.
$$

### 32. Solving for Steady-State Concentrations

Solve each equation for steady-state concentrations:

$$
\begin{aligned}
x_1 &= \frac{k_4 y_1}{k_1[S]_e + k_4}, \\
x_2 &= \frac{k_{-2}y_2 + k_1[S]_i x_1}{k_2 + k_{-1}[P]_i}, \\
y_1 &= \frac{k_{-4}x_1 + k_3[P]_e y_2}{k_4 + k_{-3}[S]_e}, \\
y_2 &= -x_1 - x_2 - y_1 + 1.
\end{aligned}
$$

$$
J = k_4 y_1 - k_{-4}x_1 = \frac{k_1k_2k_3k_4\Big([S]_i[P]_e - K_1K_2K_3K_4[S]_e[P]_i\Big)}{\text{a 16 term expression} - \text{use symbolic matlab}}.
$$
